# Enhanced conformational exploration of protein loops using a global parameterization of the backbone geometry

**DOI:** 10.1101/2022.06.21.497022

**Authors:** Timothée O’Donnell, Frédéric Cazals

## Abstract

Flexible loops are paramount to protein functions, with action modes ranging from localized dynamics contributing to the free energy of the system, to large amplitude conformational changes accounting for the repositioning whole secondary structure elements or protein domains. However, generating diverse and low energy loops remains a difficult problem.

This work introduces a novel paradigm to sample loop conformations, in the spirit of the Hit-and-Run (HAR) Markov chain Monte Carlo technique. The algorithm uses a decomposition of the loop into tripeptides, and a novel characterization of necessary conditions for Tripeptide Loop Closure to admit solutions. Denoting *m* the number of tripeptides, the algorithm works in an angular space of dimension 12m. In this space, the hyper-surfaces associated with the aforementioned necessary conditions are used to run a HAR-like sampling technique.

On classical loop cases up to 15 amino acids, our parameter free method compares favorably to previous work, generating more diverse conformational ensembles. We also report experiments on a 30 amino acids long loop, a size not processed in any previous work.

## 1 Introduction

### Protein loops

Protein loops are structural components playing various roles in protein function, as illustrated by the following examples. Enzymes typically involve conformational changes of loops for the substrate (resp. product) to enter (resp. leave) the active site [1]. Membrane transporters implement complex efflux mechanisms resorting to loops changing the relative position of (essentially) rigid domains [2]. In the humoral immune response, the binding affinity of antibodies for antigens is modulated by the dynamics of loops called complementarity determining regions (CDRs) [3]. In G-Protein-Coupled Receptors, extracellular loops binding to ligands trigger signal transduction inside the cell [4].

From the experimental standpoint, these complex phenomena are studied using structure determination methods. However, the structural diversity of loops often results in a low signal to noise ratio, yielding difficulties to report complete polypeptide chains. As a matter of fact, a recent study on structures from the PDB showed that about 83% of structures solved at a resolution of 2.0Å or worse feature missing regions, which for 90% of them are located on loops or unstructured regions [5].

### Loop modeling strategies

From the theoretical standpoint, loop mechanisms are best described in the realm of energy landscapes [6], which distinguishes between structure, thermodynamics, and dynamics. In terms of structure, one wishes to characterize active conformations and important intermediates in functional pathways. In assigning occupation probabilities to these states, one treats thermodynamics, while transitions between the states correspond to dynamics. While all atom simulations can naturally be used to explore the conformational variability of loops, their prohibitive cost prompted the development of simplified strategies, which we may ascribed to four tiers.

First, continuous geometric transformations can be used to deform loops, e.g. based on rotations of rigid backbone segments sandwiched between two *C_α_* carbons. Such methods, which include `Crankshaft` [7] and Backrub [8, 9], proved effective to reproduce motions observed in crystal structures. However, they are essentially limited to hinge like motions.

Second, a loop may be deformed using loop closure techniques solving an inverse problem which consists in finding the geometric parameters of the loop so that its endpoints obey geometric constraints. Remarkably, various such methods have been developed at the interface of structural biology and robotics [10, 11, 12, 13, 14]. Using loop closure techniques, the seminal concept of *concerned rotations* was introduced long ago to sample loop conformations [15]: first, the prerotation stage changes selected internal degrees of freedom (dof) and brakes loop connectivity; second, the postrotation step restores loop closure using a second set of dof. While early such strategies used solely dihedral angles only [15], more recent ones use a combination of valence and dihedral angles [16, 17]. The latter angles indeed provide a finer control on the amplitude of angular changes in the postrotation stage, and therefore of atomic displacements. A specific type of loop closure playing an essential role is Tripeptide Loop Closure (TLC), where the gap consists of three amino, and loop closure is obtained using the six (*φ, ψ*) angles of the three *C_α_* carbons [18, 19, 13, 20].

Third, considering a loop as a sequence of protein fragments stitched together, high resolution structures from the protein data bank (PDB) can be used to sample its conformations [21, 22]. These methods are greedy/incremental in nature, and the exponential growth of solutions results in a poorer sampling of residues in the middle of the loop. Also, they suffer from the bias inherent to the PDB structures, which favors metastable conformations. As a matter of fact, it has been shown recently using Ramachandran statistics that conformations found in the PDB are less diverse than those yielded by reconstructions in the rigid geometry model [23].

Finally, several classes of methods may be combined. For example, exploiting structural data to bias the choices of angles used to perform loop closure yields a marked improvement in prediction accuracy [24]. More recently, a method growing the two sides of a loop by greedily concatenating (perturbed) tripeptides, before closing the loop using TLC has been proposed [25].

Despite intensive research efforts, predicting large amplitude conformational changes, and/or predicting thermodynamic quantities for long loops, say beyond 12 amino acids, remains a challenge [26, 27]. These difficulties owe to the high dimensionality of loop conformational space, and also to the subtle biophysical constraints that must be obeyed.

### Contribution

This work develops a new paradigm to explore the conformational space of flexible protein loops, able to deal with loop length that were out of reach. While our method relies on the tripeptide loop closure, it is, to the best of our knowledge, the first one exploiting a global continuous parameterization of the conformational space on the loop studied. This parameterization is based on the rigidity of peptide bodies (the four atoms *C_α_* – *C* – *N* – *C_α_*), which is used to define initial conditions for the individual TLC problems and couple them.

Our presentation is organized as follows: Sec. 2 provides a high-level description of the method; Sec. 3 present experiments. Finally, Sec. 4 discusses future work.

The supplementary sections provide all mathematical, algorithmic, and in silico validation details.

## 2 Algorithm overview

### 2.1 Geometric model and ingredients

We consider a loop *L* consisting of *M* = 3 × *m* amino acids, including one or two a.a. on the boundary of the loop if necessary to obtain a multiple of three. We work in the rigid geometry model [28], in which bond lengths, valence angles, and peptide bond dihedral angle are fixed. In this model, the internal geometry of each tripeptide is defined by 12 angles coding the internal geometry of the tripeptide [18], whence an overall angular configuration space *A* of dimension 12m for the m tripeptides. (As we shall see later, this model can be relaxed, see Rmk. 10.)

Our algorithm uses a strategy similar to Hit-and-Run (HAR) [29] to sample a region 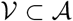 (Fig. 1). The region 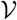 defines necessary conditions for the *m* TLC problems to admit solutions. This region is explored by shooting random curves, and intersections between the curves and the hyper-surfaces bounding 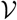 are used to generate configurations of the whole loop. Individual solutions to the *m* TLC problems are then obtained in a subset 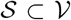. This region is further restricted to 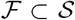, to avoid steric clashes between any pair of {*N, C_α_, C, O, C_β_*} atoms. (A steric clash is defined from the ratio between the inter atomic distance versus the sum of van der Waals radii.) The Cartesian product of solutions for the *m* tripeptides defines the new conformations of the loop *L*. We now introduce these ingredients in turn.

**Figure 1:**
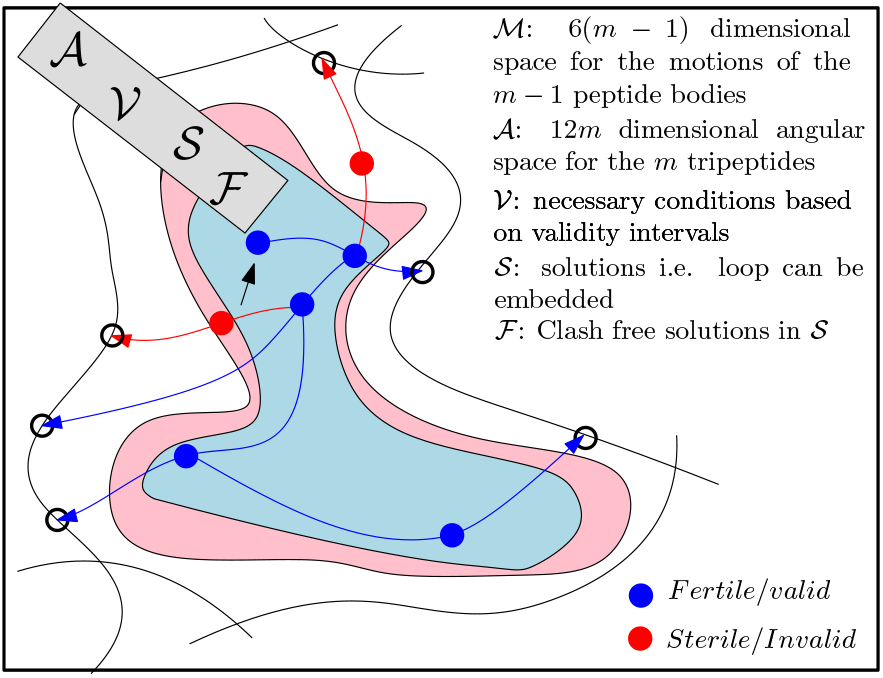
Sampling a loop involving *m* tripeptides: spaces involved, and algorithm overview. Spaces used: 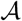: a 12m dimensional angular space coding the internal geometry of all tripeptides; 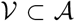: subspace characterized by necessary conditions for the *m* individual TLC problems to admit solutions; 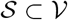: subspace such that the TLC associated with each individual tripeptide admits solutions; 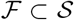: subspace such that the solutions to TLC do not yield any steric clash between any {*N, C_α_, C, O, C_β_*} atom pair. The Hit-and-Run algorithm is started at the point indicated by an arrow. It is used to find intersection (empty bullets) between 1D trajectories (blue curves) in the angular space of the tripeptides, and hyper-surfaces bounding the regions defining necessary conditions for the m individual TLC problems to admit solutions. One point is then generated on the curve segment joining the staring point and the intersection point. This point is fertile if all TLC problems admit solutions, and sterile otherwise. The number of conformations obtained is the product of the individual numbers for the *m* tripeptides. The process starts again from a fertile point with at least one solution without any steric clash.

#### Geometric model

The four atoms making up the peptide bond (*C*_*α*;1_, *C*_1_, *N*_2_, *C*_*α*;2_) form a rigid body termed the *peptide body* (Fig. S1). For the sake of exposure, we call the two segments *C*_*α*;1_ – *C*_1_ and *N*_2_ – *C*_*α*;2_ the *legs* of the tripeptide, and the tripeptide minus its legs the *tripeptide core*. We model the loop as a sequence of peptide bodies *P_k_* connecting tripeptides cores 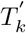 (Fig. 2):

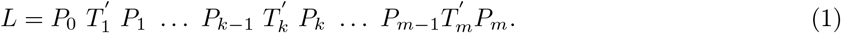

**Figure 2:**
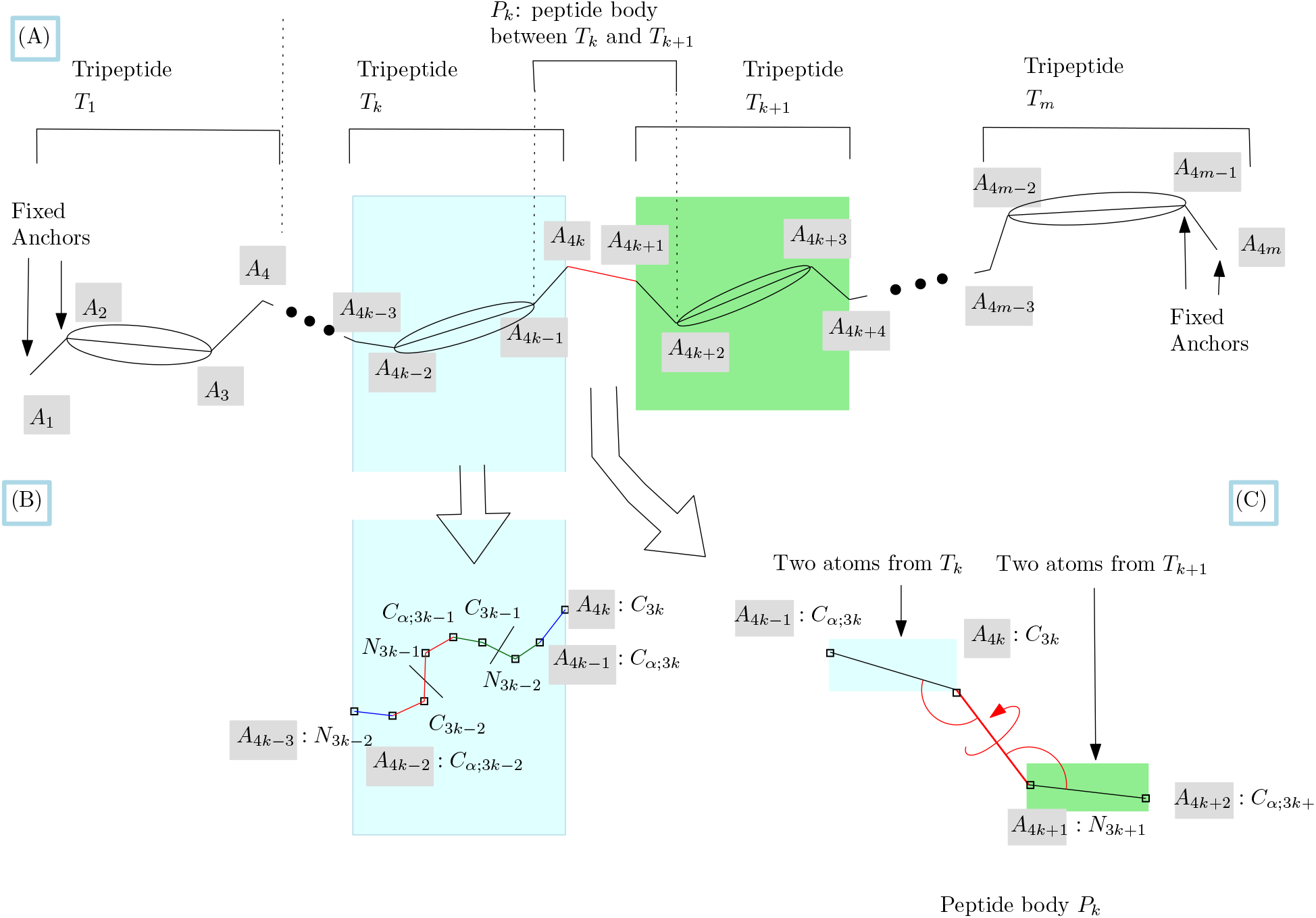
Loop decomposition into tripeptides and peptide bodies, and associated geometric model. **(A)** The *k*-th tripeptide is represented by two segments (its two legs) and an ellipsis. In red, the peptide bond between the consecutive tripeptides *T_k_* and *T*_k+1_. The peptide body encompasses the peptide bond, as well as one atom to the left and the right. **(B)** Atoms within the k-th tripeptide. **(C)** Geometry of the peptide bond linking tripeptides *T_k_* and *T*_*k*+1_ with constrained bond lengths, valence angles, and torsion angle – in red. These four atoms form the rigid body *P_k_*.

(Nb: strictly speaking, *P*_0_ and *P_m_* contain each two atoms of the loop *L*.) The main idea to generate conformations of *L* is to sample the positions of peptide bodies independently using rigid motions, and then, to solve individual TLC problems. To describe this strategy more precisely, the following ingredients are needed.

Finally, note that in Experiments, a loop refers to a set of structures with the same sequence and anchor positions (first two and last two atoms) which can be superimposed via a rigid motion.

#### Tripeptide loop closure

Tripeptide Loop Closure is a method computing all possible valid geometries of a tripeptide, under two types of constraints [18]. First, the first and last two atoms of the tripeptide, *i.e*. its legs, are fixed. Second, all internal coordinates are fixed, except the six (*φ, ψ*) dihedral angles of the three *C_α_* carbons. TLC admits at most 16 solutions corresponding to the real roots of a degree 16 polynomial. These solutions have been shown to be geometrically diverse (atoms are moving up to 5Å), so that one may use the metaphor of *teleportation* to move from one to another. Also, solutions retain a low potential energy [23]. Solving TLC can be done using three rigid bodies associated with the three edges of the triangle involving the three *C_α_* carbons. The rotations of these rigid bodies are described by three angles *τ*_1_, *τ*_2_, *τ*_3_, two of which can be eliminated to yield the degree 16 polynomial. The coefficients of this polynomial depends on 3 × 4 = 12 angles describing the internal geometry of the tripeptide [18]. This 12 dimensional space is denoted 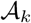 for the tripeptide *T_k_*. Taking the Cartesian product of the individual angular spaces of the *m* tripeptides yields a 12m dimensional space denoted 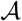.

#### Necessary conditions for TLC to admit solutions

In the angular space 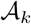, we have recently exhibited a region 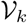 defining necessary conditions for TLC to admit solutions [30]. For a given tripeptide, this region is defined from 24 implicit equations involving the 12 variables parameterizing TLC. The corresponding space for all tripeptides is denoted 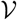. This space contains the solution space 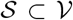, such that each tripeptide admits solutions. This space is further restricted to obtain 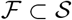 devoid of steric clash between any pair of {*N, C_α_, C, O, C_β_*} atoms.

#### Identifying active constraints with Hit-and-Run

To sample 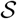, we use the Hit-and-Run (HAR) technique invented long ago to identify redundant hyperplanes in linear programs [29]. In a nutshell, given a starting point inside a polytope, HAR iteratively proceeds as follows: shoot a random ray inside the polytope and identify the nearest hyperplane intersected; generate a random point onto the segment defined by the starting and the intersection point; iterate. Since then, this algorithm has been modified to generate points following a Gaussian distribution, a key step in the computation of the volume of polytopes [31]. Other random walks serving similar purposes are billiard walk and Hamiltonian Monte Carlo [32, 33], as well as walks based on piecewise deterministic processes [34]. In the sequel, we use HAR to sample a high dimensional curved region.

### 2.2 Algorithm: wrapping up

We present two algorithms termed *unmixed* and *mixed*, coping differently with the rigidity of the peptide bond.

#### Unmixed loop sampler

Similarly to HAR, our algorithm consists of consecutive steps, called embedding steps. Each step generates a conformation *L*’ of the loop *L* by moving the peptide bodies. Given the internal coordinates of *L*’, we solve *m* individual TLC problems, one for each tripeptide, and take the Cartesian product of individual solutions.

To see how the conformation *L*’ is generated, let *SE*(3) be the special Euclidean group representing rigid motions (translation+rotation) in 3D. The *m* — 1 peptide bodies being rigid bodies, we move them in 3D space using rigid motions parameterized over the motion space 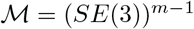. We consider a (random) ray in 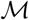, whose parameter *t* is called the *time*. Every point on this ray defines a rigid motion applied to each peptide body. Since the tripeptide legs are moving due to this motion, the 12 angular coordinates of each tripeptide become time dependent. We use the image of the ray in the angle space 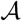 to find intersections with the hyper-surfaces defining the aforementioned necessary conditions (Fig. 1). In a manner similar to HAR, these intersections are used to generate a random point in the validity space 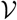. Each such point encodes an internal geometry for each tripeptide, so that TLC can be solved for each individual tripeptide. The solutions to the individual TLC problems are then combined, retaining one at random or all of them. The ability to generate efficiently points in 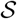 depends on the stringency of necessary conditions defining 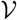, that is to say on the volume of the region 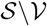.

Combining these steps yields algorithm 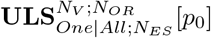, whose parameters are as follows: *One|All* a flag indicating how many solutions are retained at each embedding step, *N_ES_* the number of embedding steps, *N_V_* the number of random trajectories followed in motion space, *N_OR_* the output rate (the number of steps in-between the ones where conformations get harvested), and *p*_0_ the starting configuration.

#### Mixed loop sampler

To alleviate the constraint of fixed peptide bodies throughout the simulation, we also provide a two-step variant denoted 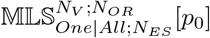. In short, every other HAR step, the loop is shortened by three residues (two a.a. on a random end, one on the other), and a HAR step is performed for this reduced model. One solution is then picked at random, and the updated positions of the peptide bodies used for the next HAR step.

## 3 Experiments

### 3.1 Material and methods

#### Implementation

Our implementation is sketched in Sec. S4.2. Consider a loop together with a valid starting point *p*_0_. First, the 12(*m* – 1) Cartesian coordinates of the peptide bodies are extracted, together with the 12 Cartesian coordinates of the two loop anchors (4 points in total). Then, the steps are iteratively performed as described above for the unmixed and mixed versions of the loop sampler.

We compare our samplers against the state-of-the-art method `MoMA-LS` [25] discussed in Introduction. We note however that the comparison cannot be done on par for three reasons. On the one hand, `MoMA-LS` samples three *ω* angles in the loop before using tripeptide loop closure. Using a distribution learned from the data may restrict the conformational space explored. On the other hand, `MoMA-LS` also samples the `ω` angle preceding the first tripeptide of the loop (Fig. S10); this degree of freedom induces a rotation of all atoms in the loop, including *C*_*α*;1_ which is fixed in our algorithm. This filtering step is subtle, since removing conformations may reduce the conformational diversity, but may also push the system further away, fostering exploration.

#### Loops tested

Several loop datasets have been assembled, see e.g. [35, 36, 26, 25]. Note that a loop refers to a set of structures with the same sequence and anchor positions which can be superimposed via a rigid motion. Most of these loops comprise between 12 and 15 amino acids. In the sequel, we focus on three such loops.

- <u>PTPN9-MEG2</u>. A 12 a.a. long loop found in the in human protein tyrosine phosphatase PTPN9-MEG2 [37, 38], between residue 466 and 477. For this case, four conformations (aka landmarks) have been crystallized: *L*_0_: 4GE2.pdb/chain A, *L*_1_: 2PA5.pdb/chain A, *L*_2_: 4GE6.pdb/chain B, *L*_3_: 4ICZ.pdb/chain A. Interestingly, three of these loops form a cluster (lRMSD < 0.1, Table S1), while *L*_3_ is significantly different (lRMSD > 1.5).
- <u>CCP-W191G</u>. A 15 a.a. long loop found in cytochrome C peroxidase (CCP), a water-soluble heme-containing enzyme reducing hydrogen peroxide (*H*_2_*O*_2_) to water. CCP contains three cavities which are hydrophobic (cavity: L99A), slightly polar (cavity: L99A/M102Q), and anionic (cavity: W191G), the latter binding almost exclusively small monocations. Out of the several crystal structures reported [39], one of them features N-methyl-1-phenylmethanamine – N-Methylbenzylamine for short in the W191G cavity. This binding is of interest, as the aforementioned 15 a.a. long loop flips out by nearly 12Å, opening the cavity to the bulk solvent for the entry/exit of the ligand [39].
- <u>CDR-H3-HIV</u>. To illustrate the ability of our method to handle long loops as a whole, we process a 30 a.a. long complementarity-determining region (CDR H3) loop, one of the longest CDR observed in human antibodies [40]. Broadly neutralizing antibodies against the human immunodeficiency virus type of 1 (HIV- 1) exhibit two typical features, namely an extensive affinity maturation (accomplished over long periods of time), and an exceptionally long heavy chain CDR.

### 3.2 Conformational diversity

To assess the conformational diversity of a set of conformations generated, we plot the root mean square fluctuations (RMSF) of the 3m heavy atoms {*N, C_α_, C*} of the loop backbone, in the form of boxplots. (Recall that the RMSF of a given atom is the stdev of distances between its positions and their center of mass.)

#### Loop PTPN9-MEG2

We first analyze the RMSF values observed for the loop PTPN9-MEG2 (Fig. 3). A general observation is the bell shape traced by the RMSF median marks, which is expected since the middle of the loop incurs less steric constraints than its endpoints. To compare the contenders, the RMSF plots for `MoMA-LS` converge rapidly. A median of ~ 2 – 3Å in the middle of the loop is obtained, with numerous extreme/outlier configurations. Our algorithm needs more steps to stabilize, reaching a stable distribution for 5000 conformations. Overall, our methods generate RMSF fluctuations larger than those from `MoMA-LS`, with 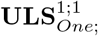. and 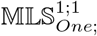. yielding median RMSF values ~ 5 – 6Å and ~ 8Å respectively near the center of the loop.

**Figure 3:**
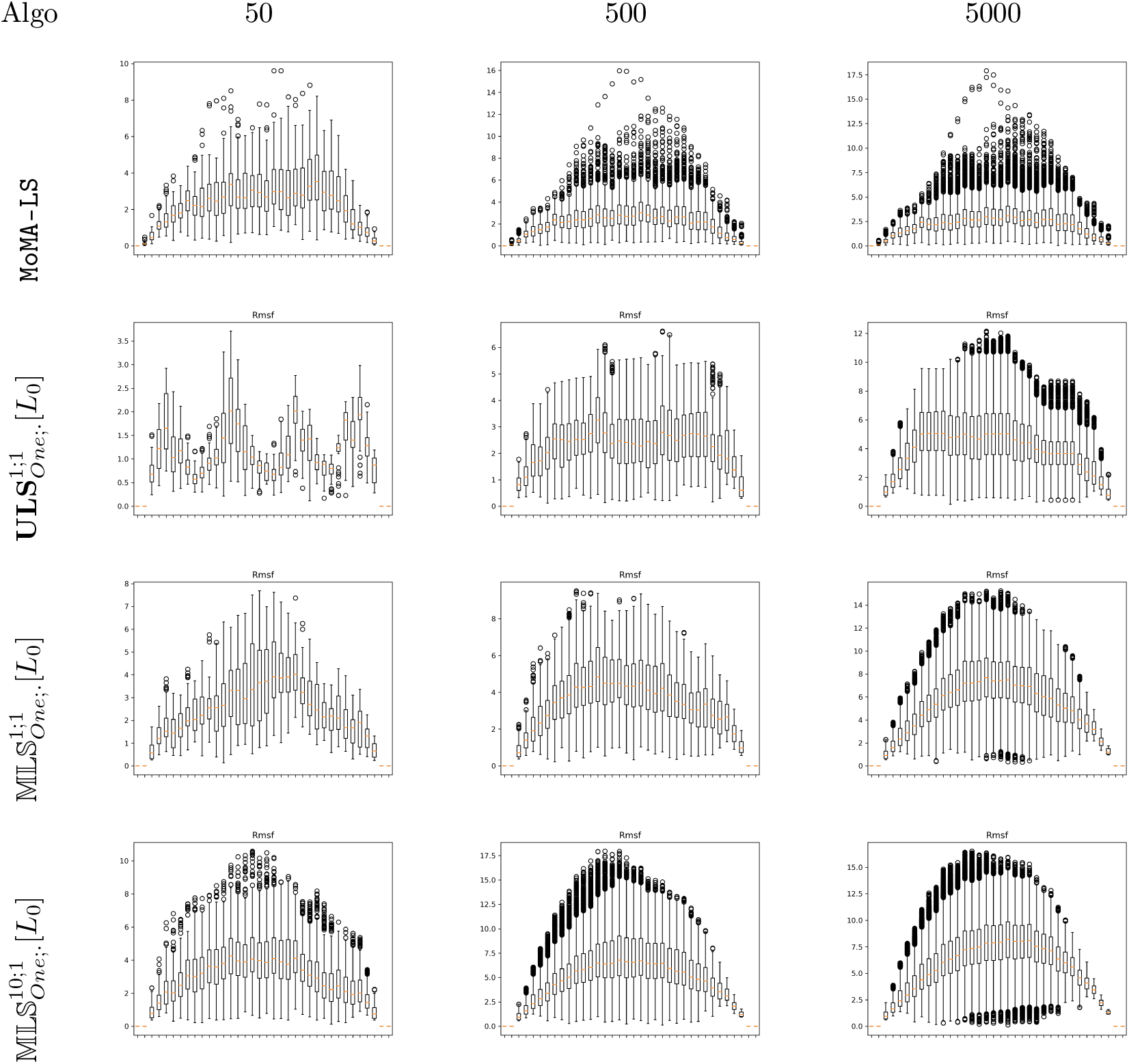
Loop PTPN9-MEG2: Backbone RMSF for the 12 amino acid long loop PTPN9- MEG2. Simulations started from the conformation/landmark *L_o_* – see text. Each tick on the x-axis corresponds to a heavy atom of the loop – 36 in this case. For `MoMA-LS`, note that only one atom is fixed on the left hand side of the loop, since the ω angle preceding the loop is also sampled.

Our plots also shed light on the various ingredients of our method. A marked difference is observed between 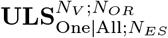 and 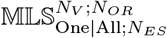. The RMSF plots of the former contain plateaus of length 4 corresponding to the atoms found in rigid peptide bodies. Those of the latter do not, a consequence of the shift shift along the backbone inherent to the removal of three amino acids.

Otherwise, an important point is the stability of our method with respect to the parameter One|All and to the number of vectors *N_V_*. Beyond 500 conformations, little variation is actually observed (Fig. 3 versus Fig. S6 and Fig. S7). (This observation can also be raised from Tables 1 and Tab. S2.). The fact that one solution provides sufficient information, is practically important since retaining all solutions per step is practically impossible for long loops.

**Table 1:** Loop PTPN9-MEG2: exploration to reach landmark conformations. Using option *One* to retain a single solution per step. Four conformations of loop PTPN9-MEG2 form two clusters: *L*_0_,*L*_1_,*L*_2_ and *L*_3_. For `MoMA-LS`, we compute min and max lRMSD distances to these landmarks. For 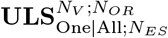 and 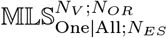, starting from *L*_0_, we compute the *max*lRMSD (resp. *min*lRMSD values) values to assess the ability to get away from the cluster (resp. approach conformation *L*_3_).

| | $L_1$ | $L_2$ | $L_3$ |
| --- | --- | --- | --- |
| | $min/max\vert IRMSD$ | $min/max\vert IRMSD$ | $min/max\vert IRMSD$ |
| MoMA-LS, 50 | 1.00/3.81 | 1.03/3.80 | 1.38/4.11 |
| MoMA-LS, 500 | 0.78/4.29 | 0.77/4.30 | 1.11/4.70 |
| MoMA-LS, 5000 | 0.74/4.92 | 0.73/4.94 | 0.99/4.97 |
| $\mathbf{ULS}_{one;50}^{1;1}[L_0]$ | 0.57/2.51 | 0.56/2.52 | 1.59/2.63 |
| $\mathbf{ULS}_{one;50}^{10;1}[L_0]$ | 0.40/3.50 | 0.40/3.50 | 1.44/3.84 |
| $\mathbf{ULS}_{one;50}^{1;1/4}[L_0]$ | 0.89/2.82 | 0.88/2.83 | 1.76/3.36 |
| $\mathbf{ULS}_{one;50}^{10;1/4}[L_0]$ | 0.52/4.49 | 0.51/4.50 | 1.45/4.99 |
| $\mathbf{ULS}_{one;500}^{1;1}[L_0]$ | 0.57/4.23 | 0.56/4.26 | 1.59/4.31 |
| $\mathbf{ULS}_{one;500}^{10;1}[L_0]$ | 0.40/4.66 | 0.40/4.68 | 1.44/5.09 |
| $\mathbf{ULS}_{one;500}^{1;1/4}[L_0]$ | 0.89/4.92 | 0.88/4.95 | 1.76/5.10 |
| $\mathbf{ULS}_{one;500}^{10;1/4}[L_0]$ | 0.52/5.23 | 0.51/5.27 | 1.45/5.52 |
| $\mathbf{ULS}_{one;5000}^{1;1}[L_0]$ | 0.57/5.27 | 0.56/5.30 | 1.59/5.43 |
| $\mathbf{ULS}_{one;5000}^{10;1}[L_0]$ | 0.40/5.35 | 0.40/5.37 | 1.44/5.86 |
| $\mathbf{ULS}_{one;5000}^{1;1/4}[L_0]$ | 0.89/5.34 | 0.88/5.36 | 1.76/5.79 |
| $\mathbf{ULS}_{one;5000}^{10;1/4}[L_0]$ | 0.52/5.35 | 0.51/5.39 | 1.45/5.80 |
| $\mathbf{MLS}_{one;50}^{1;1}[L_0]$ | 0.57/3.43 | 0.57/3.44 | 1.57/3.66 |
| $\mathbf{MLS}_{one;50}^{10;1}[L_0]$ | 0.52/4.81 | 0.51/4.83 | 1.03/5.23 |
| $\mathbf{MLS}_{one;50}^{1;1/4}[L_0]$ | 1.78/4.59 | 1.77/4.60 | 2.02/4.88 |
| $\mathbf{MLS}_{one;50}^{10;1/4}[L_0]$ | 1.32/5.22 | 1.33/5.23 | 1.38/5.87 |
| $\mathbf{MLS}_{one;500}^{1;1}[L_0]$ | 0.57/4.69 | 0.57/4.71 | 1.57/4.88 |
| $\mathbf{MLS}_{one;500}^{10;1}[L_0]$ | 0.52/5.77 | 0.51/5.79 | 1.03/6.28 |
| $\mathbf{MLS}_{one;500}^{1;1/4}[L_0]$ | 1.78/5.53 | 1.77/5.56 | 1.90/5.84 |
| $\mathbf{MLS}_{one;500}^{10;1/4}[L_0]$ | 1.32/5.75 | 1.33/5.78 | 1.38/6.29 |
| $\mathbf{MLS}_{one;5000}^{1;1}[L_0]$ | 0.57/5.67 | 0.57/5.69 | 1.57/6.26 |
| $\mathbf{MLS}_{one;5000}^{10;1}[L_0]$ | 0.52/5.93 | 0.51/5.96 | 1.03/6.50 |
| $\mathbf{MLS}_{one;5000}^{1;1/4}[L_0]$ | 1.71/5.91 | 1.71/5.95 | 1.61/6.32 |
| $\mathbf{MLS}_{one;5000}^{10;1/4}[L_0]$ | 1.32/5.93 | 1.33/5.96 | 1.38/6.50 |

#### CCP-W191G

The patterns for this slightly longer loop are similar to those observed for the previous one, so that we focus solely on the most striking point. Interestingly, despite the lack of sampling of the *ω* angle, our algorithms reach a max RMSF circa 7.5Å, while `MoMA-LS` culminates at about 3.7Å(Fig. 4 and Fig. S8).

**Figure 4:**
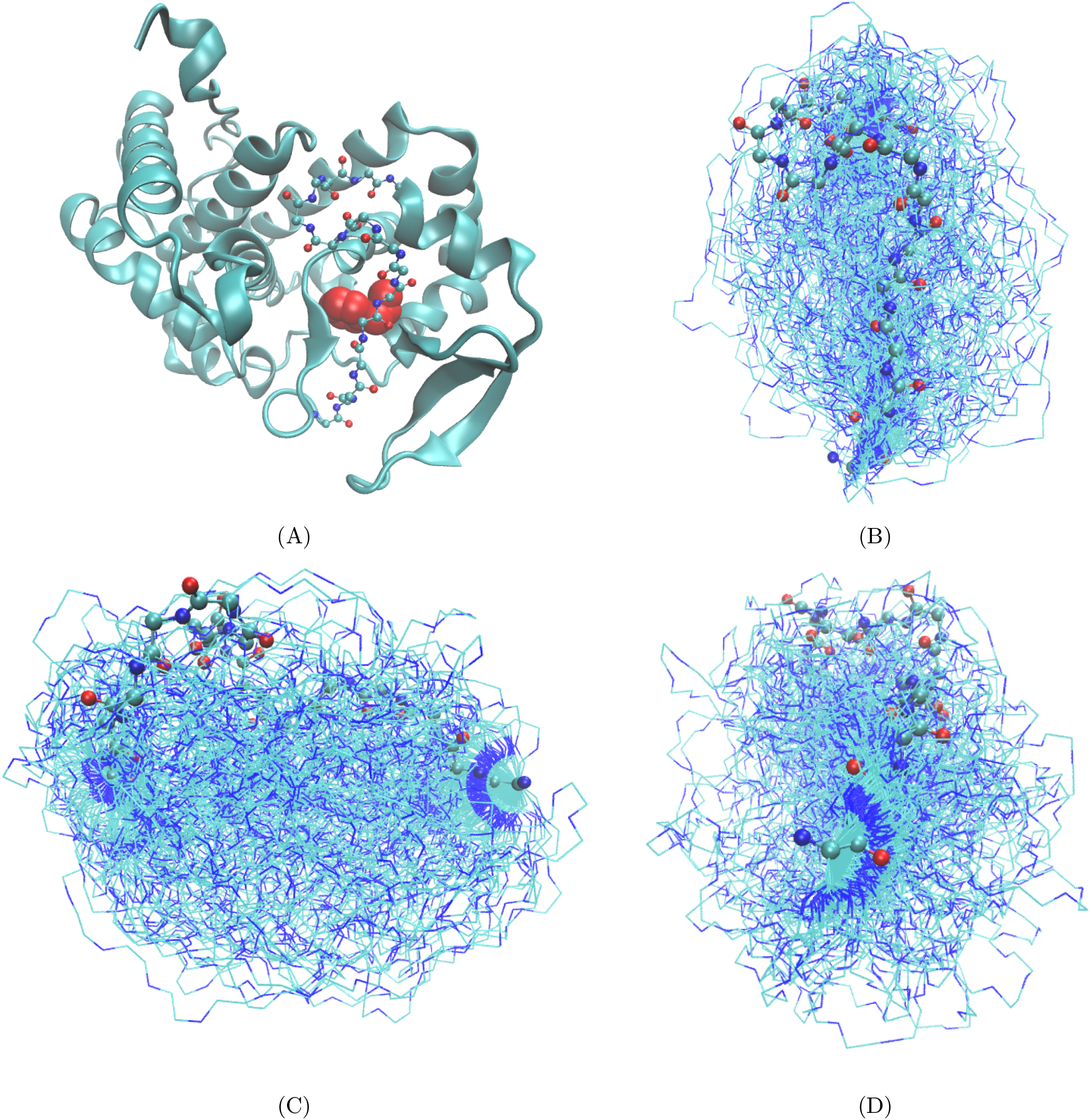
Loop CCP-W191G, 15 amino acids. Loop found in cytochrome C peroxidase (CCP). Loop specification: pdbid: 2rbt, chain X, residues 186-200. Conformations generated by algorithm 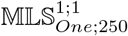. **(A)** Overview of the protein: cartoon mode: protein; CPK mode: loop; VDW representation: ligand N- Methylbenzylamine. **(B,C,D)** Top, side, front view of the loop conformations. Protein omitted for the sake of clarity.

The ability to generate such diverse ensembles is clearly an advantage over more classical methods such as Molecular Mechanics/Generalized Born Surface Area (MM/GBSA) which fail from sampling conformations as diverse as 12Å[39].

#### CDR-H3-HIV

Loops beyond 15 a.a. are usually considered to be beyond reach [26, 27]. To illustrate the capabilities of our method, we process a 30 a.a. long loop CDR-H3-HIV (Fig. 5), one of the longest CDR observed in human antibodies [40]. The CDR3 resembles an axe, with a handle and a head (Fig. 5(A)). This CDR represents alone 42% of the surface area exposed by the CDRs [40].

**Figure 5:**
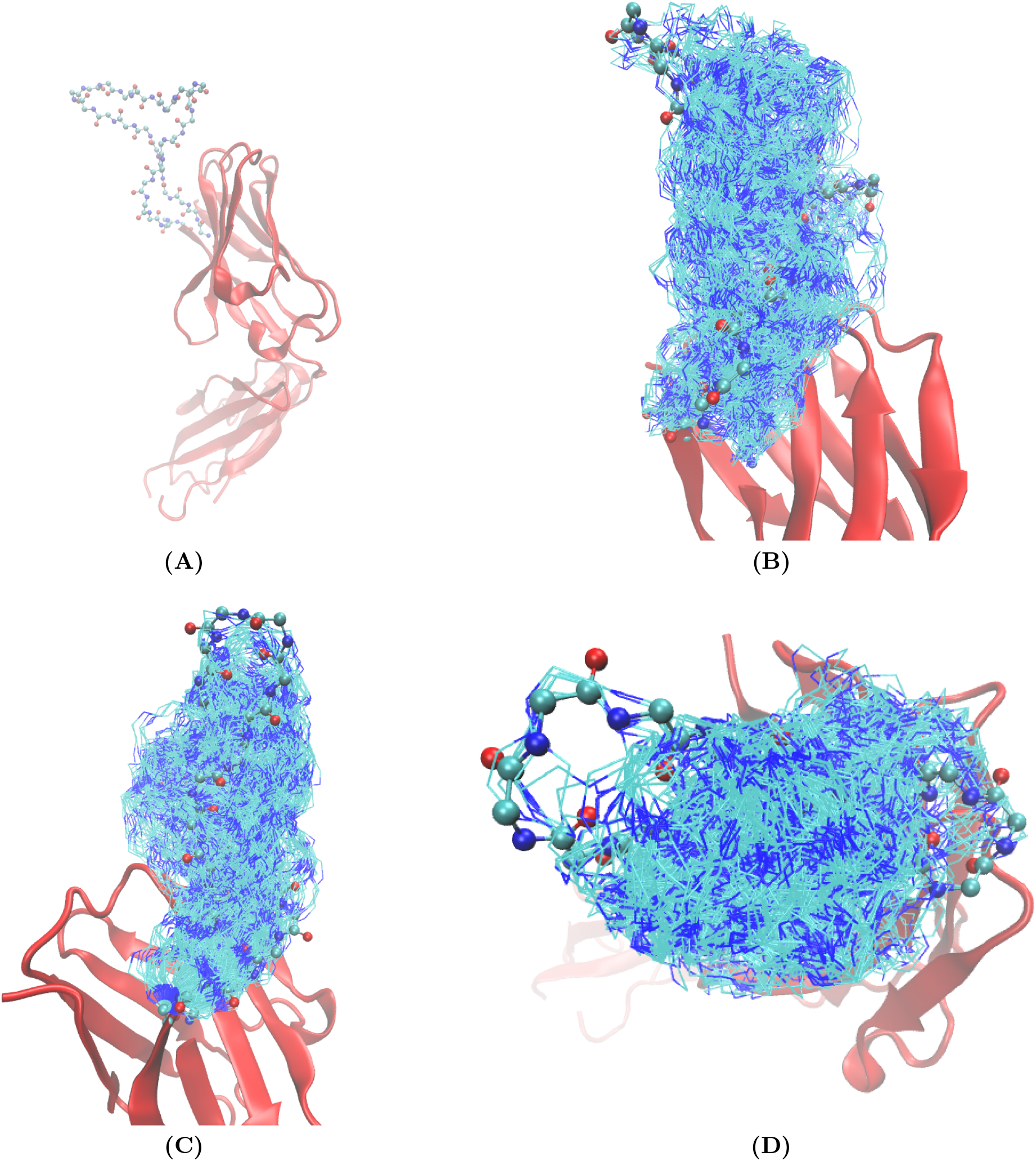
CDR-H3-HIV, 30 amino acids. The loop is a complementarity-determining region (CDR-H3) from PG16, an antibody with neutralization effect on HIV-1 [40]. Loop specification: pdbid: 3mme; chain A; residues: 93-100, 100A-100T, 101, 102. Conformations generated by algorithm 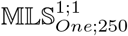. **(A)** Variable domain (red) and the 30 a.a. long CDR3. **(B,C,D)** Side/front/top view of 250 conformations.

Remarkably, compared to the two loops just discussed, `MoMA-LS` exhibits a much larger diversity (Fig. S9). Naturally, the longer the loop, the larger the benefits of also sampling the `ω` angle preceding the loop. The RMSF plots for our algorithm show a flattened bell shape curve 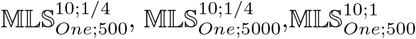 and 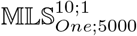, with a maximum RMSF near 12Å.

It has been speculated that the head of this CDR3 can substantially deform, possibly to maneuver into a recessed epitope [40]. Our simulations mitigates this intuition (Fig. 5(B,C,D)). On the one hand, while the *middle* of the head does deform substantially, in particular in the vertical direction, the front and the back appear quite rigid. On the other hand, the stem of the axe exhibits a substantial lateral flexibility. Naturally, these preliminary observations call for further structural analysis in the presence of the antigens.

### 3.3 Exploration of the conformational landscape

To assess the ability of the algorithm to explore a complex conformational landscape, we focus on loops for which several conformations have been obtained experimentally. Consider a set {*L_j_*},*j* = 1,…,*J* of *J* loop conformations, called *landmarks*. To assess the amount of conformational space explored, we generate conformations, and check the min and max lRMSD distances of these conformations to all landmarks.

#### Loop PTPN9-MEG2

We choose *L*_0_ as a starting point, since it is furthest away from *L*_3_. The previous pairwise distances are of special interest in the context of the 2-cluster structure of the four conformations of PTPN9-MEG2 (Table S1). In the sequel, due to the aforementioned similarity between the options *One* and *All*, we focus on the results yielded by the former.

We first observe that min lRMSD values from 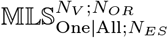 tends to be larger than those from 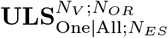.

This is expected, since the mixed sampler uses conformations of peptide bodies yielded by TLC (the aforementioned *teleportation* metaphor), as opposed to continuous moves for the unmixed sampler. Also trivially expected is a larger [min, max] lRMSD interval when more runs are performed (parameter <u>N_V_</u>).

Starting from *L*_0_, we first study the ability to move away from the cluster *L*_0_/*L*_1_/*L*_2_ (maxlRMSD values for columns *L*_1_ and *L*_2_, Table 1). For a fixed number of conformations (50/500/5000), the lRMSD observed for our algorithms are significantly larger than those obtained with the loops from `MoMA-LS`.

Also starting from *L*_0_, we next investigate the speed at which we approach the significantly different conformation *L*_3_ (minlRMSD values for column *L*_3_, Table 1). The values reported by our methods are slightly worse than those from `MoMA-LS` (Table 1): best `MoMA-LS`: 0.99Å; best 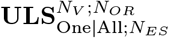: 1·47Å; best 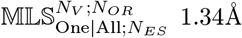. However as noticed above, `MoMA-LS` also samples the *ω* angle preceding the loop. Inspecting *ω* values, one obtains: *ω*(*L*_0_): –177°; *ω*(*L*_3_): –165°; *ω*(best from `MoMA-LS`): –167°. It is therefore the sampling of this dihedral angle which favors `MoMA-LS`.

### 3.4 Failure rate, running time and steric clashes

Algorithms 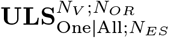 (Algorithm 2) fails as soon as one TLC does not admit any solution. This failure probability depends on the number of tripeptides, and naturally depends on the discrepancy between the two spaces 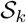 and 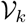, that is on the volume of the region 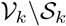. In turn, this failure naturally impacts the running time of algorithm `Loop_sampler`.

Calculations were run on a desktop DELL Precision 7920 Tower (Intel Xeon Silver 4214 CPU at 2.20GHz, 64 Go of RAM), under Linux Fedora core 32. Each HAR is processed on a single CPU core. For PTPN9- MEG2, there there is on average 0.78 failure per success when tested on 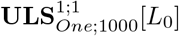 and 3.01 with 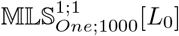.

The average time taken for one step by 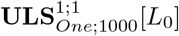 is 0.04 seconds, and 0.18 for 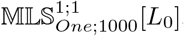. The latter algorithm involves more operations than the former, and as just noticed, also incurs a higher failure rate. Whence the increased running time.

For the long loop CDR-H3-HIV, the average failure per success becomes 2.01 for 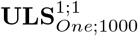 and 7.85 for 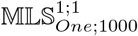. The average time per step in 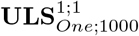 becomes 0.27 seconds, and 1.08 for 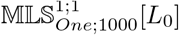.

To assess steric clashes (Sec. S3.6), we compute for a given algorithm and a set of solutions, the fraction of conformations featuring a clash. For PTPN9-MEG2, algorithms 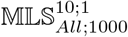 and 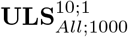 generate 20.3% and 93.5% of clashes, respectively. For CCP-W191G, algorithm 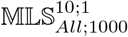 generates 10.4% of clashes. For CDR-H3-HIV and algorithm 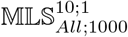, the percentage is 87.2%. As expected, steric clashes increase with the loop length, and are also more frequent when the loop resembles a hairpin.

#### Remark 1

*Parameter One/All has no impact on failure rate since all solutions are computed in any case*.

## 4 Outlook

### Method

Loops sampling methods raise difficult mathematical problems due to the high dimensionality of the parameter space, and the non linear interaction between the degrees of freedom (dof). Current state-of-the art methods belong to two main classes. The first one consists of methods relying on kinematic loop closure; such methods first perturb selected dof (the prerotation step), and proceed with loop closure (the postrotation step). However, a first difficulty is to balance the amplitude of changes incurred by pre and post dof, to avoid steric clashes during the loop closure step. Another difficulty lies in the non linear nature of the solution space. For systems involving *n* dof, such methods typically results in a solution space which is a n — 6 dimensional manifold. Sampling this manifold is usually done via back-projection upon walking the tangent space, which is numerically challenging and imposes rather local changes. A second class of methods of utmost importance exploit structures from the Protein Data Bank, and possibly resort to loop closure too. However, such methods face a combinatorial explosion when the loop length increases. As a matter of fact, modeling as a whole loops beyond 15 amino acids is still considered out of reach.

Our work introduces a new paradigm for this problem, based on a global geometric parameterization of the loop decomposed into tripeptides. The method lies in the lineage of the Hit-and-Run algorithm, invented long ago to identify redundant constraints in a linear program. Since then, HAR and related techniques have proven essential to sample high dimensional distributions in bounded and unbounded domains, yielding effective polynomial time algorithms of low complexity to compute the volume of polytopes in hundreds of dimensions [32, 31, 33, 34]. The connexion between these algorithms and loop sampling is non trivial, as using HAR to generate loop conformations involves two new ingredients. The first one is a description of the loop sampling problem in a fully dimensional conformational space, as it is the absence of codimension which removes the constraint to follow a curved manifold. We achieve such a description using the intrinsic geometric description of tripeptides. The second one is the design of necessary conditions for the individual tripeptide problems to admit solutions. These conditions can then be used in a manner akin to the hyperplanes of the polytope, to explore the region of interest and generate novel conformations.

Our results improve on those produced by a recent state-of-the-art method. On classical loop examples (12 to 15 a.a.), we show that our solutions enjoy wider RMSF fluctuations-despite not sampling ω angles. We also show that our method copes easily with a 30 a.a. long loop as a whole, a loop length usually considered beyond reach. Last but not least, it should be stressed that our method is parameter free, as the generation process does not depend on any statistical or biophysical model.

### Future work

Computational Structural Biology recently underwent a very significant progress with the advent of deep learning methods for structure prediction [41, 42]. However, such methods generally face difficulties for unstructured and/or highly flexible regions [43]. Also, they do not yield insights on the intrinsic complexity of the problem. In this context, our work opens new perspectives in structural modeling. In terms of structure, we anticipate several straightforward applications. The ability of our sampler to generate very diverse ensembles of conformations should prove key to investigate systems with highly flexible regions, including enzymes, membrane transporters, CDRs, and also intrinsically disordered proteins. The realm of thermodynamics appears more challenging. As discussed in Introduction, methods in the lineage of Conrot come with correction factors which, once incorporated into Metropolis-Hastings and Monte Carlo sampling, ensure that the correct distribution (typically canonical) is sampled. Our work primarily focuses on the geometric rather than thermodynamic setting. In fact, current sampling methods of choice are multiphase / adaptive sampling methods, including meta-dynamics, Wang-Landau, etc [44, 45]. A question of critical importance in future work will be to ensure that our exploration methods are suitable to sample NVE and/or NVT ensembles, via the calculation of densities of states (DoS). Along the way, the question of incorporating changes on internal coordinates other than dihedral angles naturally arises–but we note that changing such coordinates solely affects the conditioning of the individual TLC problems. The connexion with polytope volume calculations is a strong hint that this may indeed be the case, and that sampling micro-canonical ensembles may be possible. If so, our paradigm may eventually yield a definitive step towards effective structural and thermodynamic predictions. Meanwhile, our method can still be used in the context of global optimization and energy landscapes, which decouples structure, thermodynamics, and dynamics Upon discovering (deep) local minima, one can sample their basins [6] using classical MC methods.

## Acknowledgments

Aarushi Gupta is acknowledge for stimulating discussions. Jean-Pierre Merlet is acknowledged for providing a root finder based on ALIAS. Théo Roncalli and Viraj Agashe are acknowledged for carefully re-reading the paper.

