## Supporting Information for "Enhanced conformational exploration of protein loops using a global parameterization of the backbone geometry"

### S1 Notations: cheatsheet

**Tripeptides and the whole loop $L$.**

- Legs of a tripeptide $T_k$: the first two and last two atoms, i.e. left leg $(N_1, C_{\alpha;1})$, right leg $(C_{\alpha;3}, C_3)$. The tripeptide core $T_k^{'}$ is the tripeptide minus the legs.
- Peptide body $P_k$: the rigid body defined by the right leg and the left leg of two consecutive tripeptides.
- Loop anchors are the first two and last two atoms in the loop.
- Loop: a sequence of $m$ tripeptide:

$$L = P_0\ T_1^{'}\ P_1\ \ldots\ P_{k-1}\ T_k^{'}\ P_k\ \ldots\ P_{m-1} T_m^{'} P_m.$$

  Decomposes into left anchor + sequence of (tripeptide core+peptide body) + right anchor.

**Angular representations.**

- Tripeptide $T_k$, four tuple of angles around the $C_{\alpha;i}$: $\mathbf{A}_{k,i} = \{\alpha_{k,i}, \eta_{k,i}, \xi_{k,i-1}, \delta_{k,i-1}\}$ with $i \in \{1,2,3\}$ – counted modulo three.
- Tripeptide angular representation aggregating three four tuples: $\mathbf{A}_k = \{\mathbf{A}_{k,1}, \mathbf{A}_{k,2}, \mathbf{A}_{k,3}\}$.
- Angular conformational space of a tripeptide: 12-dimensional space $\mathcal{A}_k$
- Angular conformational space of the loop $L$: $12m$-dimensional space $\mathcal{A} = \prod_{k=1}^{m} \mathcal{A}_k$.
- Functions returning the 4 angles $\alpha, \xi, \eta$ and $\delta$ as a function of the legs of a tripeptide: $f_{(k,i)}^{(\alpha)}, f_{(k,i)}^{(\xi)}, f_{(k,i)}^{(\eta)}, f_{(k,i)}^{(\delta)}$
- Validity intervals for the angle $\tau_{k,i}$:

$$\begin{cases} \text{Initial validity interval:} I_{\tau_{k,i}} = [I_\tau^{\min}(\mathbf{A}_{k,i}), I_\tau^{\max}(\mathbf{A}_{k,i})] & \text{Sets: } \mathcal{I}_{\tau_{k,i}} = \cup I_{\tau_{k,i}} \\ \text{Rotated validity interval:} I_{\tau_{k,i}|\delta} = [I_{\tau|\delta}^{\min}(\mathbf{A}_{k,i+1}), I_{\tau|\delta}^{\max}(\mathbf{A}_{k,i+1})] & \text{Sets: } \mathcal{I}_{\tau_{k,i}|\delta} = \cup I_{\tau_{k,i}|\delta} \end{cases}$$

- Mapping from the 12 angles $\mathbf{A}_{k,i}$ into the set of validity intervals:

$$\text{DOVI}_{\tau_{k,i}}(\cdot) : \mathcal{A}_k \mapsto (\mathcal{I}_{\tau_{k,i}} \cap \mathcal{I}_{\tau_{k,i}|\delta})^4.$$

- Angular validity domain of angle $\tau_{k,i}$ for the tripeptide $T_k$: the subset of $\mathcal{A}_k$ such that $\text{DOVI}_{\tau_{k,i}}(\cdot) \neq \emptyset$.
- Depth $j$ validity intervals for $\tau_{k,i}$: $\mathcal{J}_{\tau_{k,i}}^{(j)}$.

**Motions.**

- The $6(m-1)$ dimensional space of rigid motions for the $m-1$ peptide bodies: $\mathcal{M}$
- Kinetic (dept one) validity intervals:

$$\begin{cases} I_{\tau_{k,i}}(t) = [I_\tau^{\min}(\mathbf{A}_{k,i}(t)), I_\tau^{\max}(\mathbf{A}_{k,i}(t))] \\ I_{\tau_{k,i}|\delta}(t) = [I_{\tau|\delta}^{\min}(\mathbf{A}_{k,i+1}(t)), I_{\tau|\delta}^{\max}(\mathbf{A}_{k,i+1}(t))] \end{cases}$$

**Spaces and validity domains.**

- Angular conformational space $\mathcal{A}$

$$\mathcal{A} \stackrel{Def}{=} \prod_{k=1}^{m} \mathcal{A}_k.$$

- The *angular* validity domain $\mathcal{V}$ of $L$:

$$\mathcal{V} \subset \mathcal{A} \text{ such that } \forall k, \forall i, \forall a \in \mathcal{V} : \text{DOVI}_{\tau_{k,i}}(a) \neq \emptyset.$$

- The Hit-and-Run algorithm consists of iteratively sampling a new point on $\text{Ray}_{\mathcal{V}}(p_0)V$.

- Solution space $\mathcal{S} \subset \mathcal{V}$: subspace such that TLC admits at least one solution for each tripeptide $T_k$.
- Clash free space: $\mathcal{F} \subset \mathcal{S}$: subspace such that the solutions to TLC do not yield any steric clash between any $\{N, C_\alpha, C, O, C_\beta\}$ atom pair.

### S2 Background and notations for peptides and TLC

#### S2.1 Peptides and tripeptides

**Peptides, peptide bonds, tripeptides, and protein loops.** The four atoms making up the peptide bond $(C_{\alpha;1}, C_1, N_2, C_{\alpha;2})$ form a rigid body termed the *peptide body* (Fig. S1).

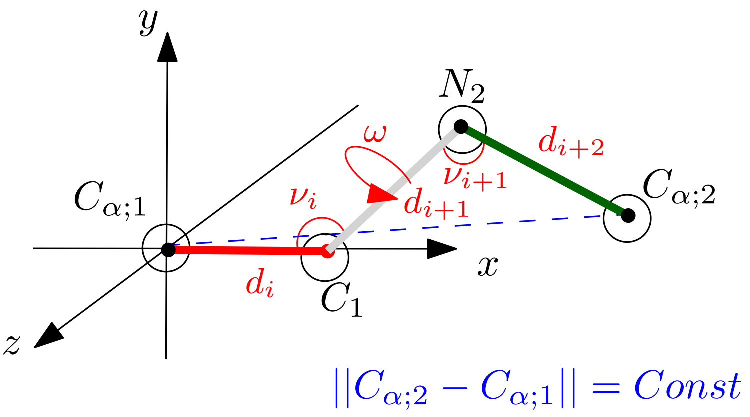


Figure S1: **The peptide body: a rigid body associated with a peptide bond.** Internal coordinates marked in red are fixed. The fixed values of the coordinates $\omega, \nu_{i+1}, d_{i+2}$ are such that the position of $C_{\alpha;2}$ is uniquely determined given positions for the previous three. Note in particular that the distance between $C_{\alpha;1}$ and $C_{\alpha;2}$ is fixed.

Atoms within the $k$-th tripeptide are denoted as $C_{\alpha;3k-2}, C_{\alpha;3k-1}, C_{\alpha;3k}$, and likewise for the $C$ and $N$ atoms (Fig. 2). We use the notation $A.$ to represent all atoms, namely $A_{4k-3} = N_{3k-2}$, $A_{4k-2} = C_{\alpha;3k-2}$, $A_{4k-1} = C_{\alpha;3k}$ and $A_{4k} = C_{3k}$.

As noticed above, the two segments $N_{3k-2}C_{\alpha;3k-2}$ and $C_{\alpha;3k}C_{3k}$ form *legs* of the tripeptide, while the tripeptide minus its legs form the *tripeptide core* $T_k^{'}$. Note that for two consecutive tripeptides, the second leg of $T_k$ and the first one of $T_{k+1}$ form the peptide bond. Note also that in the decomposition of Eq. (1), $P_0 = A_1A_2$ and $P_m = A_{4m-1}A_{4m}$ play a special role: these two fixed segments are called *anchors.*

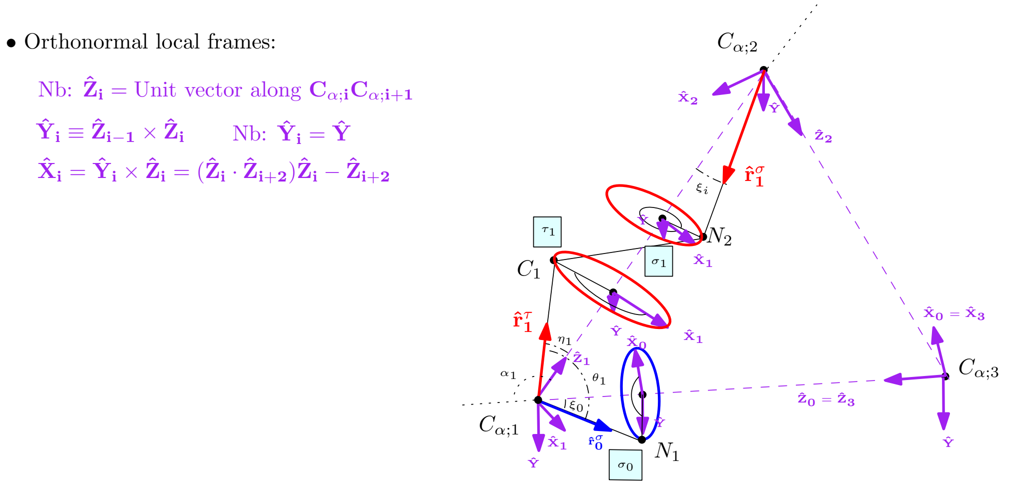


Figure S2: **Local frames and associated varibales.** Adapted from [18].

#### S2.2 Tripeptide loop closure (TLC) with fixed legs

TLC uses constraints on the tripeptide legs and internal coordinates (See Sec. 2). We may also recall that TLC induces a partition of the nine atoms in the tripeptide $T_k$ into two classes. On the one hand, the first two and the last two atoms, *i.e.* the legs, are fixed. On the other hand, the remaining five middle atoms are moving. When considering all solutions of TLC on an exhaustive database of tripeptides extracted from the PDB, these atoms move up to 5Å[23].

Solutions of TLC [18] rely on the following observations (Fig. S2, Fig. S3(A,B) [1] ):

- TLC involves three rigid bodies: the first two involve the five atoms in-between the first and third $C_\alpha$ carbons; the third one consists of the four atoms defining the legs of the tripeptide.
- The solution space of TLC can be modeled using rotation angles denoted $\{\sigma_{k,i}, \tau_{k,i}\}$ associated to the three rigid bodies. (Nb: the two angles associated with the $C_{\alpha;k,i}$ carbon are $\sigma_{k,i-1}$ and $\tau_{k,i}$.) Positions of the rigid bodies must respect the valence angles $\theta_i$ at the three $C_\alpha$ carbons. The rotation of a rigid body about its $C_\alpha - C_\alpha$ axis only impacts the valence angle constraints at its endpoints.
- Searching for solutions to the loop closure is akin to searching for rotation combinations of the angles $\{\sigma_{k,i}, \tau_{k,i}\}$ respectful of $\theta$ angles. $\sigma_{k,i-1}$ is the rotation angle of $N_i$ atoms around their corresponding axis. $\tau_{k,i}$ is the rotation angle for $C_i$ around its axis.

The geometry of the backbone can be used to define local frames at each $C_\alpha$ carbon ([18] and Fig. S3(B)), based on three vectors: $\hat{\mathbf{Z}}_{\mathbf{k,i}}$ – unit vector along two consecutive $C_\alpha$ carbons, $\hat{\mathbf{r}}^{\tau}_{\mathbf{k,i}}$ – to define the rotation of angle $\tau_{k,i}$, $\hat{\mathbf{r}}^{\sigma}_{\mathbf{k,i}}$ – to define the rotation of angle $\sigma_{k,i}$. Using these local frames, one defines the angles $\alpha_{k,i}, \xi_{k,i}, \eta_{k,i}$, with indices $i = 1, 2, 3$ – counted modulo three, for the tripeptide $T_k$ (Fig. S2):

$$\begin{cases} \alpha_{k,i} = \angle \hat{\mathbf{Z}}_{\mathbf{k,i}} \hat{\mathbf{Z}}_{\mathbf{k,i+2}}; & \alpha_{k,i} \in [0, \pi) \\ \xi_{k,i} = \angle - \hat{\mathbf{Z}}_{\mathbf{k,i}} \hat{\mathbf{r}}^{\sigma}_{\mathbf{k,i}}; & \xi_{k,i} \in [0, \pi) \\ \eta_{k,i} = \angle \hat{\mathbf{Z}}_{\mathbf{k,i}} \hat{\mathbf{r}}^{\tau}_{\mathbf{k,i}}; & \eta_{k,i} \in [0, \pi) \\ \delta_{k,i} = \angle C_{k,i} C_{\alpha;k,i}, C_{\alpha;k,i} C_{\alpha;k,i+1}, C_{\alpha;k,i+1} N_{k,i+1} & \delta_{k,i} \in [0, 2\pi) \end{cases} \tag{2}$$

**Definition. 1** *Let* $\mathbf{A}_{k,i} = \{\alpha_{k,i}, \eta_{k,i}, \xi_{k,i-1}, \delta_{k,i-1}\}$ *be the set of angles associated with* $C_{\alpha;i}$ *of the k-th tripeptide* $T_k$*. The* angular representation *of the tripeptide* $T_k$ *is the 12-tuple* $\mathbf{A}_k = \{\mathbf{A}_{k,1}, \mathbf{A}_{k,2}, \mathbf{A}_{k,3}\}$*.*

*The corresponding 12-dimensional space is denoted* $\mathcal{A}_k$*.*

#### S2.3 Tripeptide and necessary constraints for TLC

From now on, we assume that the peptide of interest is the k-th tripeptide in our loop, see Eq. (1).

In recent work [30], we have introduced necessary conditions for TLC to admit solutions. For each of the three angles $\tau_{k,i}$, these so-called *initial validity intervals* are intervals to which $\tau_{k,i}$ must belong. These intervals, which are parameterized by the angular representation of the peptide, are denoted as follows:

$$\begin{cases} \mathcal{I}_{\tau_{k,i}} = \{I_{\tau_{k,i}}\} \text{ with } I_{\tau_{k,i}} = [I^{\min}_{\tau}(\mathbf{A}_{k,i}), I^{\max}_{\tau}(\mathbf{A}_{k,i})] \\ \mathcal{I}_{\tau_{k,i}|\delta} = \{I_{\tau_{k,i}|\delta}\} \text{ with } I_{\tau_{k,i}|\delta} = [I^{\min}_{\tau|\delta}(\mathbf{A}_{k,i+1}), I^{\max}_{\tau|\delta}(\mathbf{A}_{k,i+1})] \end{cases} \tag{3}$$

There are two intervals of each type, and their pairwise intersection results in four so-called *depth one validity intervals* or DOVI. As established in [30], the bounds of these angles depend on the values

$$\arccos \frac{+\cos(\theta_i \pm \xi_{i-1}) + \cos\eta_i \cos\alpha_i}{\sin\eta_i \sin\alpha_i}. \tag{4}$$

[1] When talking of individual tripeptides $i$ is used as an index with $i \in \{1, 2, 3\}$. These indices are counted mod 3, that is $i - 1 = i + 2$.

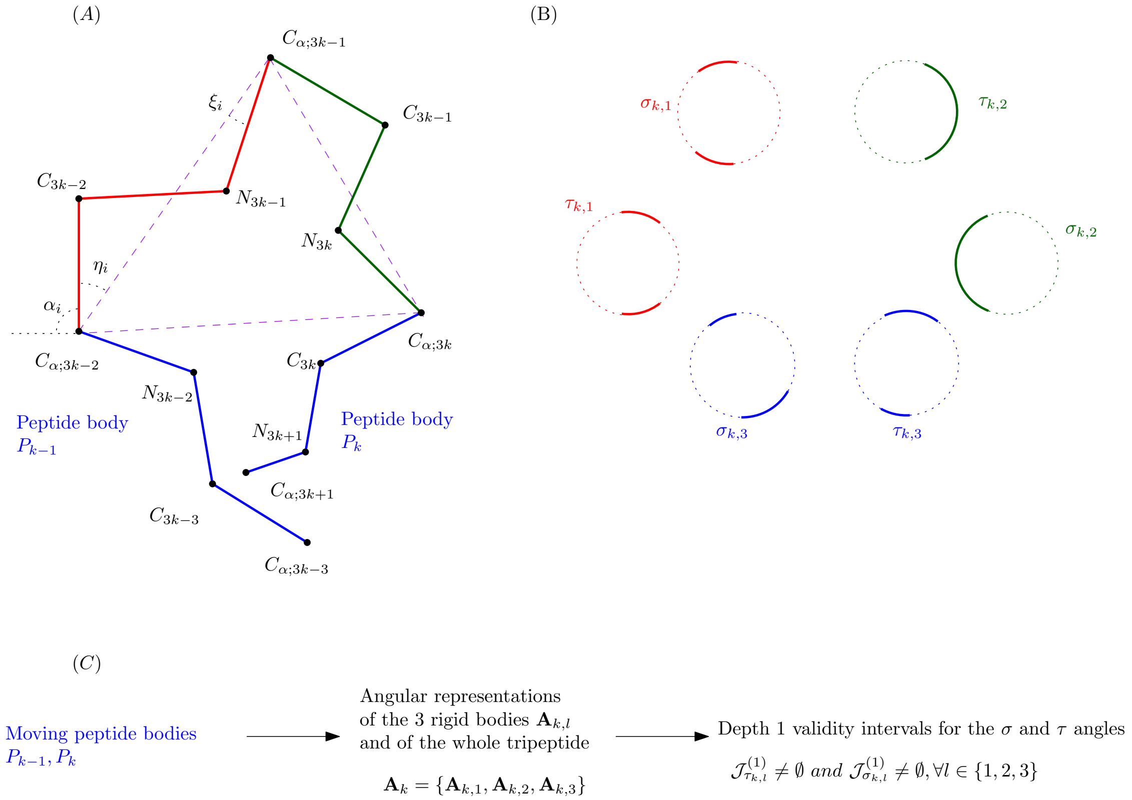


Figure S3: **Geometric model used for an individual tripeptide.** **(A)** Tripeptide with moving legs. Given internal coordinates and two rigid bodies around a tripeptide the $C_\alpha$ triangle can be defined together with $\{\alpha_i, \eta_i, \xi_i\}$ angles. **(B)** Depth one validity intervals $\mathcal{J}^{(1)}_{\sigma_i}$ and $\mathcal{J}^{(1)}_{\tau_i}$. **(C)** Illustration of the relationship between rigid body positions, $\{\alpha_i, \eta_i, \xi_i\}$ angles and the *depth one validity constraint*.

For a given tripeptide, we may consider the mapping from its angular representation in the angle space $\mathcal{A}_k$ to the validity intervals:

$$\text{DOVI}_{\tau_{k,i}}(\cdot) : \mathcal{A}_k \mapsto (\mathcal{I}_{\tau_{k,i}} \cap \mathcal{I}_{\tau_{k,i}|\delta})^4. \tag{5}$$

That is, upon fixing the angular representation of the tripeptide (Def. 1), we obtain up to four validity intervals, or the empty set if the four intersections are empty. As reported in the companion paper [30], our necessary conditions are rather tight.

**Remark 2** *The function* $DOVI_{\tau_{k,i}}$ *is obtained using the interval* $I_{\tau_{k,i}}$ *whose definition requires the angles* $\alpha_{k,i}, \eta_{k,i}, \xi_{k,i-1}$ *for* $I_{\tau_{k,i}}$*, and the interval* $I_{\tau_{k,i}|\delta}$ *whose definition requires the angles* $\alpha_{k,i+1}, \eta_{k,i+1}, \xi_{k,i}, \delta_{k,i}$*. The number of parameters is thus seven. For the sake of conciseness, we use the supersets* $\mathbf{A}_{k,i}$ *and* $\mathbf{A}_{k,i+1}$*. See [30] for details.*

### S3 Algorithm: mathematical details

#### S3.1 Tripeptides with moving legs

**Moving peptides bodies.** When considering the decomposition of Eq. (1), the $m-1$ peptide bodies move independently. The motion of one peptide body is parameterized by the special Euclidean group $SE(3)$, which combines one translation and one rotation (Fig. S4). To be more specific, let $S^2$ be the sphere of directions in $\mathbb{R}^3$, and $A$ a positive real number. The motion space $\mathcal{R}$ for one peptide body is defined via the motion space

$$\mathcal{R} : (S^2 \times [0, A)) \times (S^2 \times [0, 1/A)) \subset SE(3). \tag{6}$$

The term $S^2 \times [0, A)$ codes the translation defined by a unit vector and a real number in $[0, A)$, while the term $S^2 \times [0, 1/A)$ codes the rotation defined by an angle about a direction given by a unit vector on $S^2$ and a real number in $[0, 1/A)$. Therefore, specifying a random rigid motion for each peptide body requires $2(m-1)$ unit vectors. We pool these vectors into a $6(m-1)$-dimensional vector denoted $V$ in the sequel. Each rigid body is simultaneously translated along the first unit vector and rotated around the second using a corresponding kinematic function. The value of $A$ defines the speed of translation in such a function relative to the rotation speed, *i.e.* if $A = 0.5$ then $1/A = 2$ and the corresponding rigid body will rotate four times as fast as it translates. We use the default value $A = 1$, as we hardly noticed any incidence for this parameter (data not shown). Summarizing, the overall motion space for peptide bodies is the $6(m-1)$ dimensional space:

$$\mathcal{M} = \mathcal{R}^{m-1}. \tag{7}$$

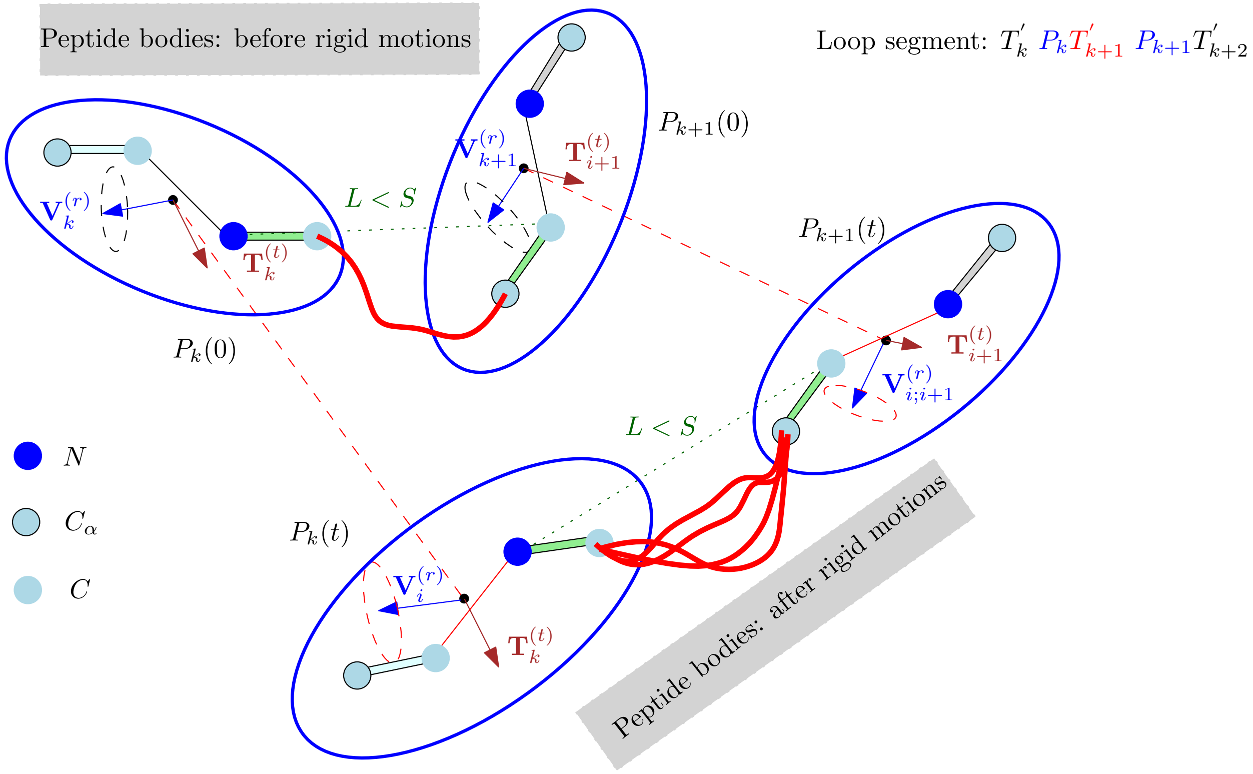


Figure S4: **Interpolation in the space of rigid motions $\mathcal{R}$ and associated transformations applied to rigid bodies.** The figure features two peptide bodies $P_k$ and $P_{k+1}$ in the loop segment $T_k^{'}$ $P_k T_{k+1}^{'}$ $P_{k+1} T_{k+2}^{'}$. The initial positions of the bodies are denoted $P_k(0)$ and $P_{k+1}(0)$ respectively; these bodies must satisfy a distance constraint materialized by the green line segment – *length* $< S$. Each rigid body undergoes a translation (unit vectors $\mathbf{T}_k^{(t)}$ and $\mathbf{T}_{k+1}^{(t)}$ respectively) composed with a rotation (unit vectors $\mathbf{V}_k^{(r)}$ and $\mathbf{V}_{k+1}^{(r)}$ respectively). The positions corresponding to time $t$ are denoted $P_k(t)$ and $P_{k+1}(t)$ respectively. The distance between the last $C_\alpha$ of $P_k(t)$ and the first $C_\alpha$ of $P_{k+1}(t)$ is constrained by the triangular inequality (Sec. S3.2.6). This constraint is represented by the maximum length $S$ on the figure.

**Remark 3** *By the Mozzi–Chasles' theorem, our rigid motion can be modeled as a screw motion. The corresponding analytical form is used to find intersections with the surfaces defining the TLC necessary constraints.*

#### S3.2 Sampling rigid motions for a single peptide body

##### S3.2.1 Using a 1-parameter family in the motion space

We restrict motions in $\mathcal{M}$ to a a 1-parameter family, performing the following linear interpolation defined by vector $V$:

$$\text{Ray}(V) = \{\gamma(t) = Id + tV, \text{ with } \gamma(0) = Id\}. \tag{8}$$

The restriction of this one parameter family to each peptide body defines a rigid transformation

$$\gamma_k : [0,1] \mapsto SE(3), \gamma_k(0) = Id, \tag{9}$$

such that the position of the k-th peptide body $P_k(t)$ at time $t$ satisfies

$$P_k(t) = \gamma_k(t) P_k(0). \tag{10}$$

We now explicit the analytical expression for $P_k(t)$.

##### S3.2.2 Rigid body representation

We have noticed that the two anchor points on each side of a peptide bond (four atoms in total), form a rigid body (Fig. 2(C)). This rigid body enjoys three translational and three rotational degrees of freedom (dof).

Note that using homogeneous coordinates, the matrix $4 \times 4$ matrix giving the coordinates of this rigid body reads as

$$P_k = \begin{pmatrix} A_{4i-1} & A_{4i} & A_{4i+1} & A_{4i+2} \\ 1 & 1 & 1 & 1 \end{pmatrix} \tag{11}$$

In the sequel, we will consider a parameterized such matrix, denoted $P_k(t)$, with $t$ a real number.

##### S3.2.3 Translation

- $\mathbf{U}_i^{(t)}, i = 1, \ldots, m-1$: unit vectors drawn uniformly at random on the sphere of directions on $S^2$. Used to define the directions of translations.
- $C_i^{(t)}, i = 1, \ldots, m-1$: $m-1$ uniformly random variables in $(0, 2\pi)$, to define the norm of translation vectors.
- Translation vector: $\mathbf{T}_i^{(t)} = C_i^{(t)} * \mathbf{U}_i^{(t)}, i = 1, \ldots, m-1$

Using homogeneous coordinates, the corresponding transformation reads as follows:

$$\tilde{\mathbf{T}}_{\mathbf{i}}(t) = \begin{pmatrix} 1 & 0 & 0 & t\mathbf{T}_{i;x}^{(t)} \\ 0 & 1 & 0 & t\mathbf{T}_{i;y}^{(t)} \\ 0 & 0 & 1 & t\mathbf{T}_{i;z}^{(t)} \\ 0 & 0 & 0 & 1 \end{pmatrix} = \begin{pmatrix} I & t\mathbf{T}_i^{(t)} \\ 0 & 1 \end{pmatrix} \tag{12}$$

##### S3.2.4 Rotation

- $\mathbf{U}_i^{(r)}, i = 1, \ldots, m-1$: units vectors drawn uniformly at random on the sphere of directions on $S^2$. Used to define the rotation axis.
- $C_i^{(r)}, i = 1, \ldots, m-1$: $m-1$ uniformly random variables in $(0, 2\pi)$, to define the rotation angles.
- Rotation: of angles $C_i^{(r)}$ around the direction $\mathbf{U}_i^{(r)}$.

In homogeneous coordinates:

For $\tilde{\mathbf{R}}_\mathbf{i}(t)$ consider $\theta = tC_i^{(r)}$ and $R(\theta, \mathbf{U}_i^{(r)})$ the rotation matrix corresponding to a rotation of $\theta$ around axis $\mathbf{U}_i^{(r)}$:

$$\tilde{\mathbf{R}}_\mathbf{i}(t) =$$
$$\begin{pmatrix} cos\theta + \mathbf{U}_{i;x}^{(r)2}(1-cos\theta) & \mathbf{U}_{i;x}^{(r)}\mathbf{U}_{i;y}^{(r)}(1-cos\theta) - \mathbf{U}_{i;z}^{(r)}sin(\theta) & \mathbf{U}_{i;x}^{(r)}\mathbf{U}_{i;z}^{(r)}(1-cos\theta) - \mathbf{U}_{i;y}^{(r)}sin(\theta) & 0 \\ \mathbf{U}_{i;y}^{(r)}\mathbf{U}_{i;x}^{(r)}(1-cos\theta) + \mathbf{U}_{i;z}^{(r)}sin(\theta) & cos\theta + \mathbf{U}_{i;y}^{(r)2}(1-cos\theta) & \mathbf{U}_{i;y}^{(r)}\mathbf{U}_{i;z}^{(r)}(1-cos\theta) - \mathbf{U}_{i;x}^{(r)}sin(\theta) & 0 \\ \mathbf{U}_{i;z}^{(r)}\mathbf{U}_{i;x}^{(r)}(1-cos\theta) + \mathbf{U}_{i;y}^{(r)}sin(\theta) & \mathbf{U}_{i;z}^{(r)}\mathbf{U}_{i;y}^{(r)}(1-cos\theta) - \mathbf{U}_{i;x}^{(r)}sin\theta & cos\theta + \mathbf{U}_{i;z}^{(r)2}(1-cos\theta) & 0 \\ 0 & 0 & 0 & 1 \end{pmatrix}$$
$$\tilde{\mathbf{R}}_\mathbf{i}(t) = \begin{pmatrix} (R(\theta, \mathbf{U}_i^{(r)})) & 0 \\ 0 & 1 \end{pmatrix} \tag{13}$$

The three types of constraints are equivalent as they are applied on the distance between two points. In all cases the two points are each part of a rigid body and in each cases each point will be subjected to a different translation and rotation. The kinematic function can then be expressed using the rigid body transformations.

##### S3.2.5 Complete transformation

- $P_k(0)$ is the homogeneous 3 dimension coordinates of a rigid body without translation and rotation.
- $P_k(t)$ is the homogeneous 3 dimension coordinates of a rigid body with translated and rotated using $\tilde{\mathbf{T}}_\mathbf{i}(t)$ and $\tilde{\mathbf{R}}_\mathbf{i}(t)$.
- $T_i^0$ is the translation of the center of mass of $P_k(0)$ to the origin and $T_i^{-1}$ the opposite.

Still using homogeneous coordinates we obtain:

$$\gamma_k(t) = \tilde{\mathbf{T}}_\mathbf{i}(t)T_i^{-1}\tilde{\mathbf{R}}_\mathbf{i}(t)T_i^0 \tag{14}$$

The complete transformation applied to $P_k$ becomes:

$$P_k(t) = \gamma_k(t)P_k \tag{15}$$

$\gamma_k(t)$ can be applied to any individual atom in $P_k$.

##### S3.2.6 Numerical root finding and tmax

When numerically searching for a solution to Eq. 25, an initial search interval is needed:

Given leg positions for a given tripeptide the three $C_\alpha$ carbons within satisfy a triangle inequality (S1). Using the proper indices, this constraint reads as

$$\|C_{\alpha;3i} - C_{\alpha;3i-2}\| < L_{C_{\alpha;3i-2}C_{\alpha;3i-1}} + L_{C_{\alpha;3i-1}C_{\alpha;3i}} \tag{16}$$

As noticed above, the distances With $L_{C_{\alpha;3i-2}C_{\alpha;3i-1}}$ and $L_{C_{\alpha;3i-1}C_{\alpha;3i}}$ are fixed. If this is not satisfied we have a forbidden sample as it belongs to $\mathcal{A}\backslash\mathcal{V}$.

- The triangular inequality is used to find an upper bound for numerical root finding.
- **Remark 4** *So long as the translation vectors are not the same there will always be a points where the distance is greater than a given value (Fig. S4).*
- Let $c_i$ and $c_{i+1}$ be the centers of the rotation circles for both atoms on which the constraint applies. These correspond to the orthogonal projections on their respective rotational axes.
- With $S = L_{C_{\alpha;3i-2}C_{\alpha;3i-1}} + L_{C_{\alpha;3i-1}C_{\alpha;3i}}$.
- With $r_i$ and $r_{i+1}$ the respective radii of said circles, the triangular inequality will necessarilly be invalid:

$$\left\|\tilde{\mathbf{T}}_{\mathbf{i+1}}(t)c_{i+1}(t) - \tilde{\mathbf{T}}_{\mathbf{i}}(t)c_i(t)\right\| = S + r1 + r2 \tag{17}$$

- This corresponds to a univariate second degree polynomial with one positive and one negative root. The upper limit of our initial constraint with both rotation and translation is the positive root.

In the loop the smallest of such values among all tripeptides is selected as an initial upper bound for $t_max$.

#### S3.3 Validity domain and overall configuration space $\mathcal{A}$

We now wish to use the depth one validity constraints for the $m$ peptides, whose legs are moving as just explained. To this end, we concatenate the angular representations of the $m$ tripeptides (Def. 1), and define:

**Definition. 2** *(Angular conformational space $\mathcal{A}$) The* angular conformational space *of the loop L is the* $12m$ *dimensional space defined by the product of the m angular space of the individual tripeptides:*

$$\mathcal{A} \stackrel{Def}{=} \prod_{k=1}^{m} \mathcal{A}_k. \tag{18}$$

Fixing the positions of the peptide bodies in Eq. (1) yields the angular representations of the $m$ tripeptides. We therefore define a mapping from the motion space into the global angular space:

$$f_{\mathcal{M}\to\mathcal{A}} : \mathcal{M} \mapsto \mathcal{A} \tag{19}$$

Having discussed the depth one validity interval for one tripeptide– see Eq. (5), we can finally aggregate such conditions:

**Definition. 3** *(Angular validity domain $\mathcal{V}$.) The* angular *validity domain $\mathcal{V}_k$ of the angle $\tau_{k,i}$ of the k-th tripeptide is the subset of $\mathcal{A}_k$ such that $DOVI_{\tau_{k,i}}(\cdot) \neq \emptyset$.*

*The* angular *validity domain of the loop L is the subset $\mathcal{V} \subset \mathcal{A}$ such that*

$$\forall k = 1, \ldots, m, \forall i = 1, \ldots, 3, \forall a \in \mathcal{V} : DOVI_{\tau_{k,i}}(a) \neq \emptyset.$$

Note that there are $3m$ individual angular validity domains since each tripeptide has 3 angles $\tau$.

Points in $\mathcal{V}$ satisfy necessary conditions. However, for a point $p \in \mathcal{V}$, one or several tripeptide may not admit any valid geometry. We therefore define:

**Definition. 4** *(Solution space $\mathcal{S}$) The* solution space *$\mathcal{S} \subset \mathcal{V}$ of the loop L is the subspace of $\mathcal{A}$ such that TLC admits at least one solution for each tripeptide. A point in $\mathcal{S}$ (resp. $\mathcal{V}\backslash\mathcal{S}$) is termed* fertile *(resp. sterile).*

Let $s_k$ the number of solutions yielded by TLC for a point $p \in \mathcal{S}$. The Cartesian product of these sets yields a total number of embeddings, *i.e.* conformations, equal to $\Pi_{k=1,\ldots,m} s_k$.

**Remark 5** *Note that the degrees of freedom are defined for rigid bodies in-between tripeptides while the constraints are defined within the tripeptides (Fig. 2).*

#### S3.4 Kinetic validity intervals

We now wish to use our 1-parameter family of motions to explore the solutions space $\mathcal{S}$ via an exploration of the valid space $\mathcal{V}$.

The tripeptide legs move according to the motion imposed to the peptide bodies (Eq. (10)). It is therefore possible to define a time dependent (aka kinetic) version of the angles $\mathbf{A}_{k,i}$:

$$\mathbf{A}_{k,i}(t) = (f^{(\alpha)}_{(k,i)}(t), f^{(\xi)}_{(k,i)}(t), f^{(\eta)}_{(k,i)}(t), f^{(\delta)}_{(k,i)}(t)), \tag{20}$$

with

$$\begin{cases} f^{(\alpha)}_{(k,i)}(t) & : \text{ function computing the angle } \alpha_{k,i} \text{ at time } t \\ f^{(\xi)}_{(k,i)}(t) & : \text{ function computing the angle } \xi_{k,i} \text{ at time } t \\ f^{(\eta)}_{(k,i)}(t) & : \text{ function computing the angle } \eta_{k,i} \text{ at time } t \\ f^{(\delta)}_{(k,i)}(t) & : \text{ function computing the angle } \delta_{k,i} \text{ at time } t \end{cases} \tag{21}$$

Once plugged into the intervals of Eq. (3), these functions make it possible to define a kinetic version of the four static validity intervals:

**Definition. 5** *(Kinetic validity intervals) The* kinetic validity intervals *for a given angle $\tau_{k,i}$ of a tripeptide $T_k$ are the validity intervals obtained for the time varying angles $\mathbf{A}_{k,i}(t)$:*

$$\begin{cases} I_{\tau_{k,i}}(t) = [I^{min}_{\tau}(\mathbf{A}_{k,i}(t)), I^{max}_{\tau}(\mathbf{A}_{k,i}(t))] \\ I_{\tau_{k,i}|\delta}(t) = [I^{min}_{\tau|\delta}(\mathbf{A}_{k,i+1}(t)), I^{max}_{\tau|\delta}(\mathbf{A}_{k,i+1}(t))] \end{cases} \tag{22}$$

**Remark 6** *The time dependent angles are computed as follows (Fig. S2):*

- *The fixed internal coordinates within each tripeptide are sufficient to determine the value of $\eta_{k,1}, \xi_{k,1}, \eta_{k,2}$ and $\xi_{k,2}$. (Note that these are defined in the rigid bodies associated with $C_{\alpha;3k-2}C_{\alpha;3k-1}$ or $C_{\alpha;3k-1}C_{\alpha;3k}$.)*
- *The position of the* legs *are sufficient to define $\eta_{k,3}$ and $\xi_{k,3}$.*
- *The leg positions together with the fixed internal coordinates are sufficient to compute all three $\alpha_{k,i}, i \in \{1, 2, 3\}$ angles as these angles are defined by the $C_\alpha$ triangle.*

**Remark 7** *The motions of consecutive rigid bodies is constrained by the triangle inequality between the three consecutive $C_\alpha$ atoms (Fig. S3). Indeed, these atoms must satisfy the following triangle inequality:*

$$\|C_{\alpha;3k-2}C_{\alpha;3k}\| \leq \|C_{\alpha;3k-2}C_{\alpha;3k-1}\| + \|C_{\alpha;3k-1}C_{\alpha;3k}\|. \tag{23}$$

*Note that following the rigidity of peptide bodies, the two right hand side distances are fixed.*

#### S3.5 Sampling: one step

**Sampling $\mathcal{V}$ with Hit-and-Run.** We sample the validity domain $\mathcal{V}$ using the Hit-and-Run algorithm (Fig. 1 and [29]). For a ray $\text{Ray}(V)$ in the motion space (Eq. (8)), consider the restriction of this ray to the valid space $\mathcal{V}$, that is

$$\text{Ray}_{\mathcal{V}}(V) = \{\gamma(t) \in \text{Ray}(V) \mid f_{\mathcal{M}\to\mathcal{A}}(\gamma(t)) \in \mathcal{V}\}. \tag{24}$$

The Hit-and-Run algorithm consists of iteratively sampling a new point on $\text{Ray}_{\mathcal{V}}(V)$, so that the restriction of the ray to the valid space $\mathcal{V}$ must be computed.

To see how, consider two kinetic intervals $I_{\tau_{k,i}}(t) \in \mathcal{I}_{\tau_{k,i}}$ and $I_{\tau_{k,i}|\delta}(t) \in \mathcal{I}_{\tau_{k,i}|\delta}$ as specified in Eq. (22). For these intervals, consider the limit conditions (Fig. S5):

$$\begin{cases} I^{\max}_{\tau}(\mathbf{A}_{k,i}(t)) = I^{\min}_{\tau|\delta}(\mathbf{A}_{k,i+1}(t)), \\ \text{ or } I^{\min}_{\tau}(\mathbf{A}_{k,i}(t)) = I^{\max}_{\tau|\delta}(\mathbf{A}_{k,i+1}(t)) \end{cases} \tag{25}$$

For a given $\tau_{k,i}$ angle, there are 8 such conditions, namely two (Eqs. (25)) for each of the depth one validity interval. And since there are three $\tau_{k,i}$ angles per tripeptide, we obtain 24 conditions.

With these ingredients, our algorithm operates as follows:

- Generate a random ray $\text{Ray}(V)$ in the motion space $\mathcal{M}$.
- (`get_tau_tmax`, Algorithm 1 and Sec. S3.2.6) For a given $\tau_{k,i}$ angle, find out the largest interval $[0, t_{\max}]$ such that the $\text{DOVI}_{\tau_{k,i}}$ is different from the $\emptyset$ on this interval (Nb: an upper bound on $t_{\max}$ is obtained from the triangle inequality applied to the $C_\alpha$ carbons, see remark 7.)
- (`LS_one_step`, Algorithm 2) Take the intersection of all such intervals for the $3m$ angles, generate a $t$ value on the resulting interval, and apply the corresponding motions to the tripeptide legs. This yields a candidate conformation $L_{\text{cand.}} \in \mathcal{V}$ of the loop $L$.
- (`Loop_sampler`, Algorithm 3). Test if $L_{\text{cand.}} \in \mathcal{S}$. Then, iterate `LS_one_step` until a point $L_{\text{cand.}} \in \mathcal{F}$ is obtained. Once obtained, start again from $L_{\text{cand.}}$ and iterate.

**Remark 8** *In algorithm* `LS_one_step`*, it should be noted that taking the intersection ensures that all conditions hold. But one may have $DOVI_{\tau_{k,i}} \neq \emptyset$ on other segments defined by intersection between the ray and the 24 hyper-surfaces. In practice, preliminary tests did not show a significant improvement in tracking such segments.*

**Remark 9** *In the* real random access memory model *(real RAM), which assumes exact calculations with real numbers, Algorithm* `LS_one_step` *is exact. In practice, our implementation uses multiprecision numbers and root finding routines provided by Maple [46]. Due to the cost of such operations, algorithm 2 can be further optimized, see algorithm 5.*

*Leaving the realm of multiprecision, an approximate version has also been developed to strike a compromise between exactness and performances, see* `LS_one_step_approx` *(Algorithm 4). This variant performs a regular sampling of the ray, from which $t_{max}$ is estimated.* `LS_one_step_approx` *is the version used in the experiments thereafter.*

#### S3.6 Sampling: combining several steps

We use the building block `Loop_sampler` to define two algorithms. In our Experiments, the loops assessed are those generated by these two algorithms, without any relaxation/energy minimization or post-processing.

**Unmixed loop sampler.** Combining steps of `Loop_sampler` yields algorithms $\mathbf{ULS}^{N_V;N_{OR}}_{One|All;N_{ES}}[p_0]$, whose parameters are as follows:

1. $p_0$: the starting point/conformation in space $\mathcal{S}$.
2. One|All: a point in the solution space $\mathcal{S}$ generates a total of $N_m = \Pi_{k=1,\ldots,m} s_k$ loop conformations, with $s_k$ the number of TLC solutions for the tripeptide $T_k$. The flag One|All states whether we choose one embedding at random, or keep them all.
3. $N_{ES}$: for a given HAR trajectory, the number of embedding steps performed.
4. $N_V$: number of HAR trajectories started at $p_0$, each defined by a random vector defining a ray in the motion space $\mathcal{M}$.
5. $N_{OR}$: the output rate in the form $1/n$, with $n$ the number of HAR steps performed along a HAR trajectory, before an *embedding step* is performed–as dictated by the flag One|All. An output rate of one means that all embeddings steps are exploited.

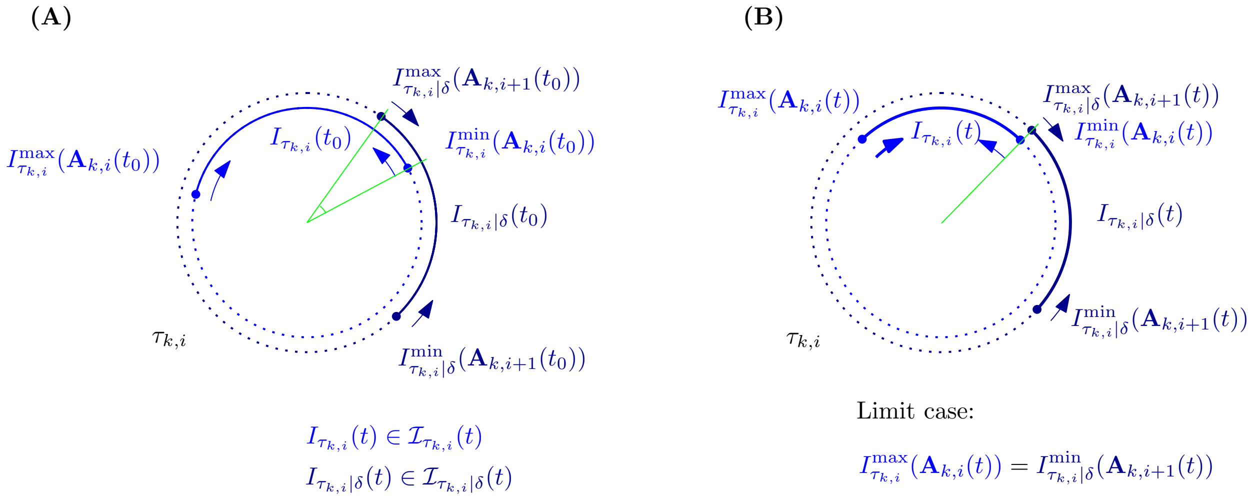


Figure S5: **Kinetic validity intervals.** We focus on a given interval pair $I_{\tau_{k,i}} \in \mathcal{I}_{\tau_{k,i}}$ and $I_{\tau_{k,i}|\delta} \in \mathcal{I}_{\tau_{k,i}|\delta}$ for the angle $\tau_{k,i}$ from tripeptide $T_k$. The legs of $T_k$ are moving with $P_{k-1}$ and $P_k$. These movements impact the positions of the interval endpoints via the angles $\mathbf{A}_{k,i}(t)$ and $\mathbf{A}_{k,i+1}(t)$. **(A)** The interiors of the two intervals intersect. **(B)** The intervals intersect on their boundary–a limit case. The arrow indicate the derivative of the endpoints of intervals with respect to time.

For example, $\mathbf{ULS}^{5;1/4}_{One;1000}$ uses five HAR trajectories with an output rate of 1/4, and 1000 embedding steps, each retaining a single embedding. Thus, the number of loop conformations returned is exactly 1250. On the other hand, $\mathbf{ULS}^{1;1}_{All;1000}$ uses a single HAR trajectory of 1000 steps with an output rate of one, retaining all solutions at each step. The number of loop conformations generated is at least 2000, and at most $1000 \times 16^m$.

**Mixed loop sampler.** In the previous version of the algorithm, peptide bodies remain rigid during the whole simulation. The two-step variant of the algorithm $\mathbb{MLS}^{N_V;N_{OR}}_{One|All;N_{ES}}[p_0]$ removes this limitation. Every other HAR step, three residues are removed from the loop endpoints (two a.a. on one end, one on the other, at random), and a HAR step is performed for this reduced model. One solution is then picked at random, and the updated positions of the peptide bodies used for the next HAR step.

**Steric clashes and collision checking.** A general post-processing strategy in loop generation consists of checking the absence of steric clash. Denoting $R_i$ and $R_j$ the van der Waals radii of two atoms $i$ and $j$, the usual criterion consists of checking that $d_{ij}/(R_i + R_j) \geq r_{min}$, usually taken in the range $0.5 - 0.7$, see [20, 25]. Upon generating a conformation, we perform this check for all pairs of $N, C_\alpha, C, O, C_\beta$ atoms in the loop. (The positions of atoms $O$ and $C_\beta$ are computed using the positions of the surrounding $N, C_\alpha, C$ atoms and fixed internal coordinates, see *e.g.* [**?**].)

**Remark 10** *We have recalled above the two types of constraints used by TLC: the legs' positions and internal coordinates. Practically, we use equilibrium values for fixed internal coordinates found in the ff14sb force field from* `AMBER`*. These internal coordinates can be changed and sampled in the course of the algorithm, an option not used in our experiments.*

*In using these standard coordinates, we assume that all tripeptides of the loop have angular parameters* $\mathbf{A}_k \in \mathcal{S}$.

### S4 Algorithms and implementation details

#### S4.1 Pseudo-code

**Algorithm 1** `get_tau_tmax`**.** For a given angle $\tau_{k,i}$, find the largest value of $t_{\max}$ of $t$ such that $\text{DOVI}_{\tau_{k,i}}(p(t)) \neq \emptyset$ on the segment $[0, t^{\Delta}_{max}]$.

**for** $I_{\tau_{k,i}}(t) \in \mathcal{I}_{\tau_{k,i}}(t)$ **do**
  **for** $I_{\tau_{k,i}|\delta}(t) \in \mathcal{I}_{\tau_{k,i}|\delta}(t)$ **do**
    $S = S \cup$ numerical solutions for Eqs. 25 $t \in [0, t^{\Delta}_{max}]$
Sort $S$ by ascending order
Let $t_l$ be the $l$-th element of $S$
$l = 1$
$u_l := \frac{t_l + t_{l+1}}{2}$
// Stop when no validity interval can be defined for $\tau_{k,i}$
**while** $\text{DOVI}_{\tau_{k,i}}(f_{\mathcal{M}\to\mathcal{A}}\gamma(u_l)) \neq \emptyset$ **do**
  $t_{max} = u_l$
  $l = l + 1$
**return** $\{t_{max}\}$

**Algorithm 2** `LS_one_step`. Given a starting point $p_0 \in \mathcal{S}$ and a random direction $V$ in the motion space $\mathcal{M}$, the algorithm finds the nearest intersection $p_{near}$ of the image of the ray $\text{Ray}(V)$ (by the map $f_{\mathcal{M}\to\mathcal{A}}$) with a surface constraint, and generates a random value on the segment $[0, t_{\max}]$. Then, applies the corresponding motion to peptide bodies of the loop $L$.

Input: $p_0 \in \mathcal{S}$: starting point in the fertile space
Input: $V$: direction in motion space
Output: a point $p_{out} \in \mathcal{V}$
Var $t^{\Delta}_{\max}$: initialized using the smallest value of $t > 0$ breaking triangular inequality in a given tripeptide
$V$: Random direction (Eq. (8))
$S = \{t^{\Delta}_{\max}\}$
**for** k $\in \{1, ..., m\}$ **do**
  **for** i $\in \{1, 2, 3\}$ **do**
    $S = S \cup$ get_tau_tmax$(\tau_{k,i})$
// Get the smallest value – most stringent condition
$t_{\max} = \min S$
// Output the next sample
$t_s \leftarrow \text{Uniform}(0, t_{\max})$
Apply the rigid transforms defined by $t_s$ to the $m - 1$ peptide bodies
**return** Loop $L$ with moved peptide bodies

**Algorithm 3** `Loop_sampler`. Given a starting point $p_0 \in \mathcal{S}$, algorithm `Loop_sampler` iterates `LS_one_step` until $L_{\text{cand.}}$ yields solution(s) for all tripeptides in the loop. This process is then repeated iteratively from $L_{\text{cand.}}$.

Input: $p_0 \in \mathcal{V}$
$p_{tmp} = p_0$
$Sample = \emptyset$
**while** not done **do**
    $is\_in\_F = false$
    **while** not $is\_in\_F$ **do**
        $is\_in\_S = false$
        **while** not $is\_in\_S$ **do**
            Generate random direction $V$
            $L_{\text{cand.}} \leftarrow$ LS_one_step($p_{tmp}, V$)
            Solve individual TLC for the $m$ peptide bodies
            **if** all $m$ tripeptide have at least one solution **then**
                $is\_in\_S = true$
        **if** $\exists$ at least one conformation with no steric clash **then**
            $is\_in\_F = True$
    $p_{tmp} = L_{\text{cand.}}$
    // Combine the individual solutions obtained for the individual tripeptides
    **if** Stop condition met **then**
        done=true

**Algorithm 4** `LS_one_step_approx`. Given a starting point $p_0 \in \mathcal{S}$, a random direction $V$ in the motion space $\mathcal{M}$, and a number of iteration $X$, the algorithm uniformly samples between 0 and $t_{\max}$, and finds the largest value $u_l$ such that $\mathrm{DOVI}_{\tau_{k,i}}(f_{\mathcal{M}\to\mathcal{A}}(\gamma(u_l))) \neq \emptyset$. It then iterates between the step were it stopped and the one before it until $\mathrm{DOVI}_{\tau_{k,i}}(f_{\mathcal{M}\to\mathcal{A}}\gamma(u_l)) \neq \emptyset$. If $X \to \infty$ the $t_{\max}$ obtained using this algorithm corresponds to the one obtained using `LS_one_step`.

Input: $p_0 \in \mathcal{S}$: starting point in the fertile space
Input: $V$: direction in motion space
Input: $X$: max number of iteration to obtain approximate solution
Output: a point $p_{out} \in \mathcal{V}$
Var $t^{\Delta}_{\max}$: initialized using the smallest value of $t > 0$ breaking triangular inequality in a given tripeptide
$V$: Random direction (Eq. (8))
$u_l := 0$
$x = 1$
// Identify the first iteration failing the condition
in_validity_space=true
**while** in_validity_space **do**
    $u_l = (x/X)t_{\max}$
    **for** $k \in \{1..m\}$ **do**
        **for** $i \in \{1,2,3\}$ **do**
            **if** $\mathrm{DOVI}_{\tau_{k,i}}(f_{\mathcal{M}\to\mathcal{A}}\gamma(u_l)) = \emptyset$ **then**
                in_validity_space=false
    $x = x + 1$
// Slice the failing interval into X bits and iterate
$t_{min} = (x-1)/Xt_{\max}$
$x = 1$
in_validity_space=true
**while** in_validity_space **do**
    $u_l = t_{min} + \frac{t_{\max}(x)}{X^2}$
    **for** $k \in \{1..m\}$ **do**
        **for** $i \in \{1,2,3\}$ **do**
            **if** $\mathrm{DOVI}_{\tau_{k,i}}(f_{\mathcal{M}\to\mathcal{A}}\gamma(u_l)) = \emptyset$ **then**
                in_validity_space=false
    $x = x + 1$
$t_{\max} = t_{min} + \frac{t_{\max}(x-1)}{X^2}$
// Output the next sample
$t_s \leftarrow \mathrm{Uniform}(0, t_{\max})$
Apply the rigid transforms defined by $t_s$ to the $m-1$ peptide bodies
**return** Loop $L$ with moved peptide bodies

**Algorithm 5 `LS_one_step`: optimized version**. In this optimized version of Algorithm 2, the upper bound $t_{\max}$ is updated incrementally for all $\tau_{k,i}$ angles, which makes it possible to seek individual roots (for a given $\tau_{k,i}$ angle) on a shorter interval.

Input: $p_{in} \in \mathcal{S}$: starting point in the fertile space
Input: $V$: direction in motion space
Output: a point $p_{out} = \mathcal{V}$

Var $t_{\max}$: initialized using the smallest value of $t > 0$ breaking triangular inequality in a given tripeptide

$V$: Random direction (Eq. (8))
**for** k $\in \{1, ..., m\}$ **do**
  **for** i $\in \{1, 2, 3\}$ **do**
    // Angle $\tau_{k,i}$: process the (at most) 24 equations
    $S = \{t_{\max}\}$
    // Process all interval pairs
    **for** $I_{\tau_{k,i}}(t) \in \mathcal{I}_{\tau_{k,i}}(t)$ **do**
      **for** $I_{\tau_{k,i}|\delta}(t) \in \mathcal{I}_{\tau_{k,i}|\delta}(t)$ **do**
        $S_{tmp} \leftarrow$ numerical solutions for Eqs. 25 $t \in [0, t_{\max}]$
        $S = S \cup S_{tmp}$
    Sort $S$ by ascending order
    Let $t_l$ be the $l$-th element of $S$
    $u_l := \frac{t_l + t_{l+1}}{2}$
    $l = 1$
    // Stop when no validity interval can be defined for $\tau_{k,i}$
    **while** $\text{DOVI}_{\tau_{k,i}}(\tau_{k,i}(u_l)) \neq \emptyset$ **do**
      $t_{\max} = t_k$
      $l = l + 1$
// Output the next sample
$t_s \leftarrow \text{Uniform}(0, t_{\max})$
Apply the rigid transforms defined by $t_s$ to the $m - 1$ peptide bodies

##### S4.2 Implementation

- Loop_sampler. The sampler generates the necessary random directions and applies the rigid transformation to each $P_k$ at each step.
- LS_tripep_validity_domain. The individual tripeptide validity domain class contain methods mapping $P_{k-1}$ and $P_k$ to $\mathcal{A}$ as well as computing $t_{\max}$ (Algo. 1)
- LS_bb_embeder. The backbone embedder: performs TLC on all tripeptides using standard internal coordinates and double precision.

#### S5 Results

##### S5.1 loops used

###### S5.1.1 `MoMA-LS` parameters

Here we summarize the parameters used in `MoMA-LS` for our experiments:

- The ratio of van der Waals radii used for collision detection is 0.5. The minimum value is used as we do not implement collision detection;
- Residue-dependent pseudo-atoms at the $C_\beta$ positions are not used for collision detection for the same reason;

| | $L_0$ | $L_1$ | $L_2$ | $L_3$ |
|---|---|---|---|---|
| $L_0$ | | 0.099 | 0.072 | 1.574 |
| $L_1$ | | | 0.087 | 1.550 |
| $L_2$ | | | | 1.559 |

Table S1: **Least RMSD matrix between landmark pairs for the loop PTPN9-MEG2.** The first three conformations form a cluster.

- Side chains are omitted as they are not considered in our algorithms as of now;
- One solution is kept for inverse kinematics as we mostly compare to the version using one solution for inverse kinematics in our algorithm;
- The number of sampled states is 50 500, or 5000.

#### S5.2 Comparisons

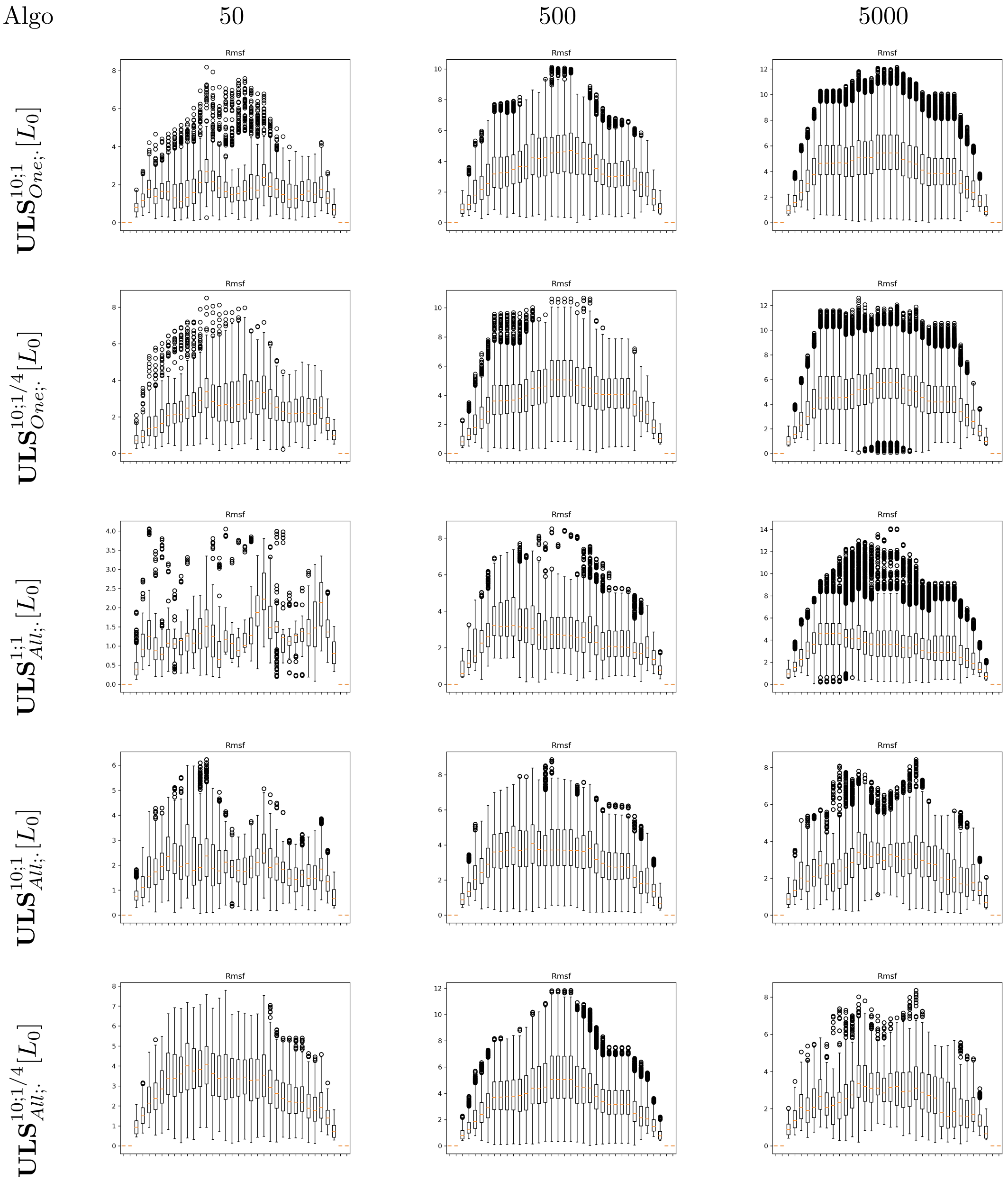


Figure S6: **Loop PTPN9-MEG2: tests with algorithm $\mathbf{ULS}^{N_V;N_{OR}}_{All;N_{ES}}[L_0]$.** Compare against Fig. 3 to see the incidence of option All.

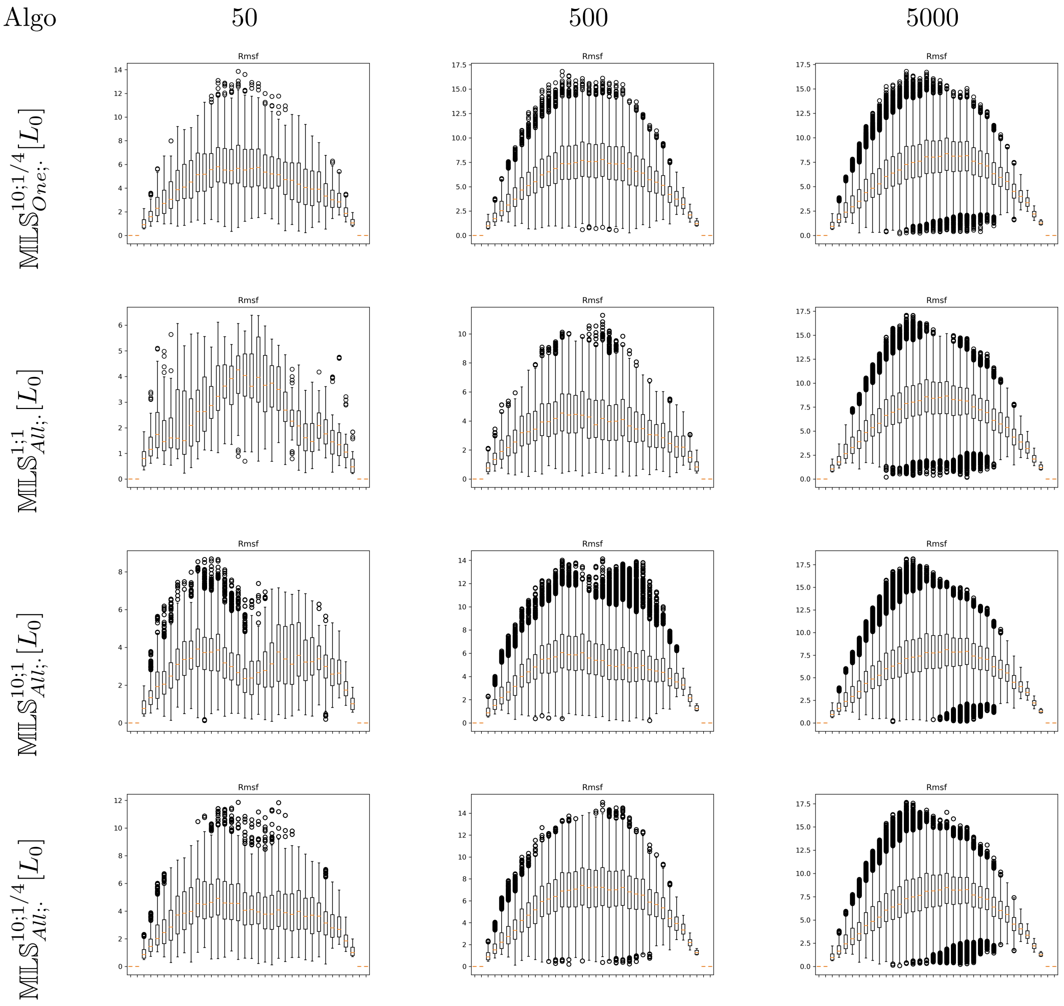


Figure S7: **Loop PTPN9-MEG2: tests with algorithm** $\mathbb{MLS}^{N_V;N_{OR}}_{All;N_{ES}}[L_0]$**.** Compare against Fig. 3 to see the incidence of option All.

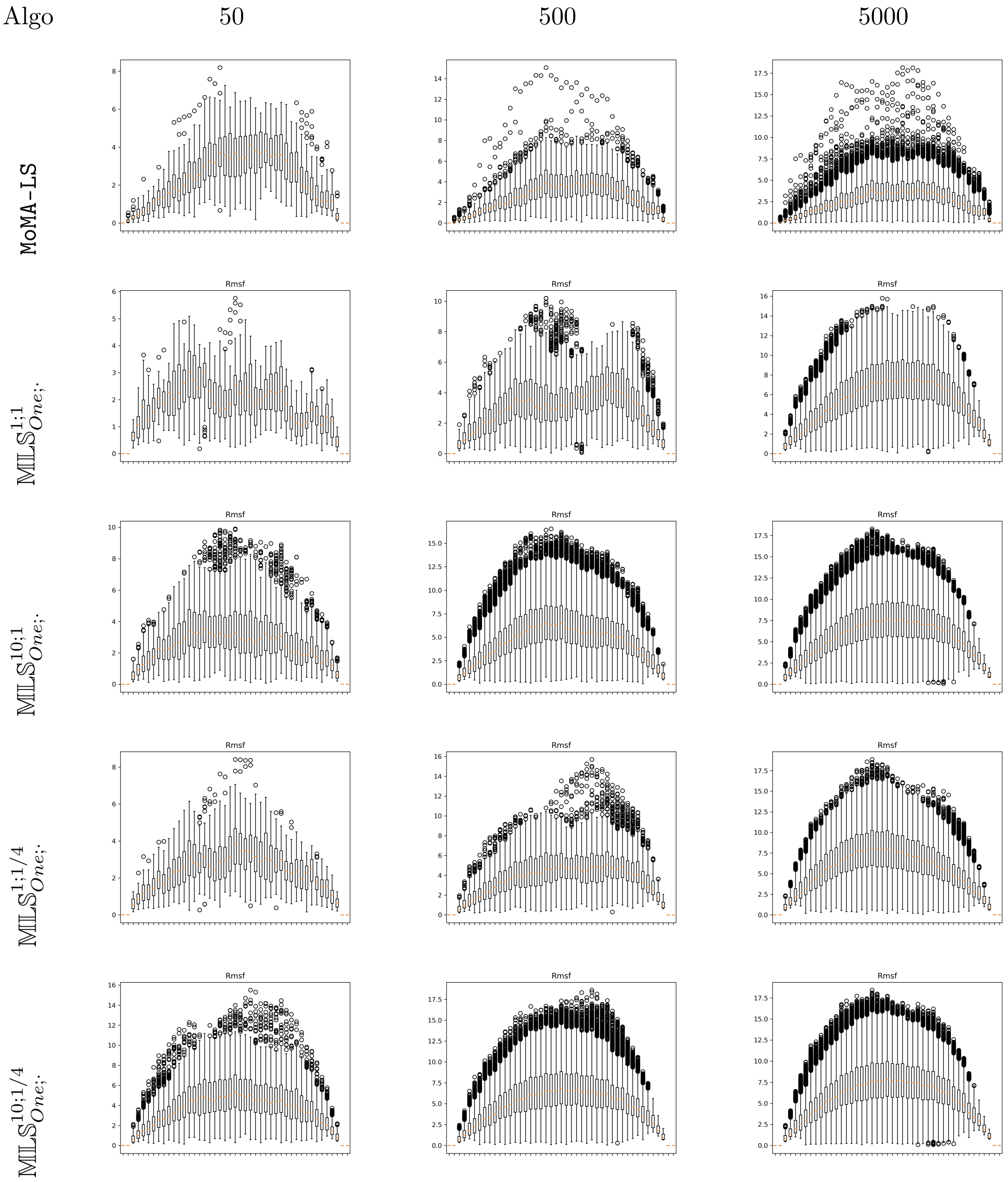


Figure S8: **Loop CCP-W191G: tests with algorithm $\mathbb{MLS}^{N_V;N_{OR}}_{one;N_{ES}}$ and MoMA-LS.**

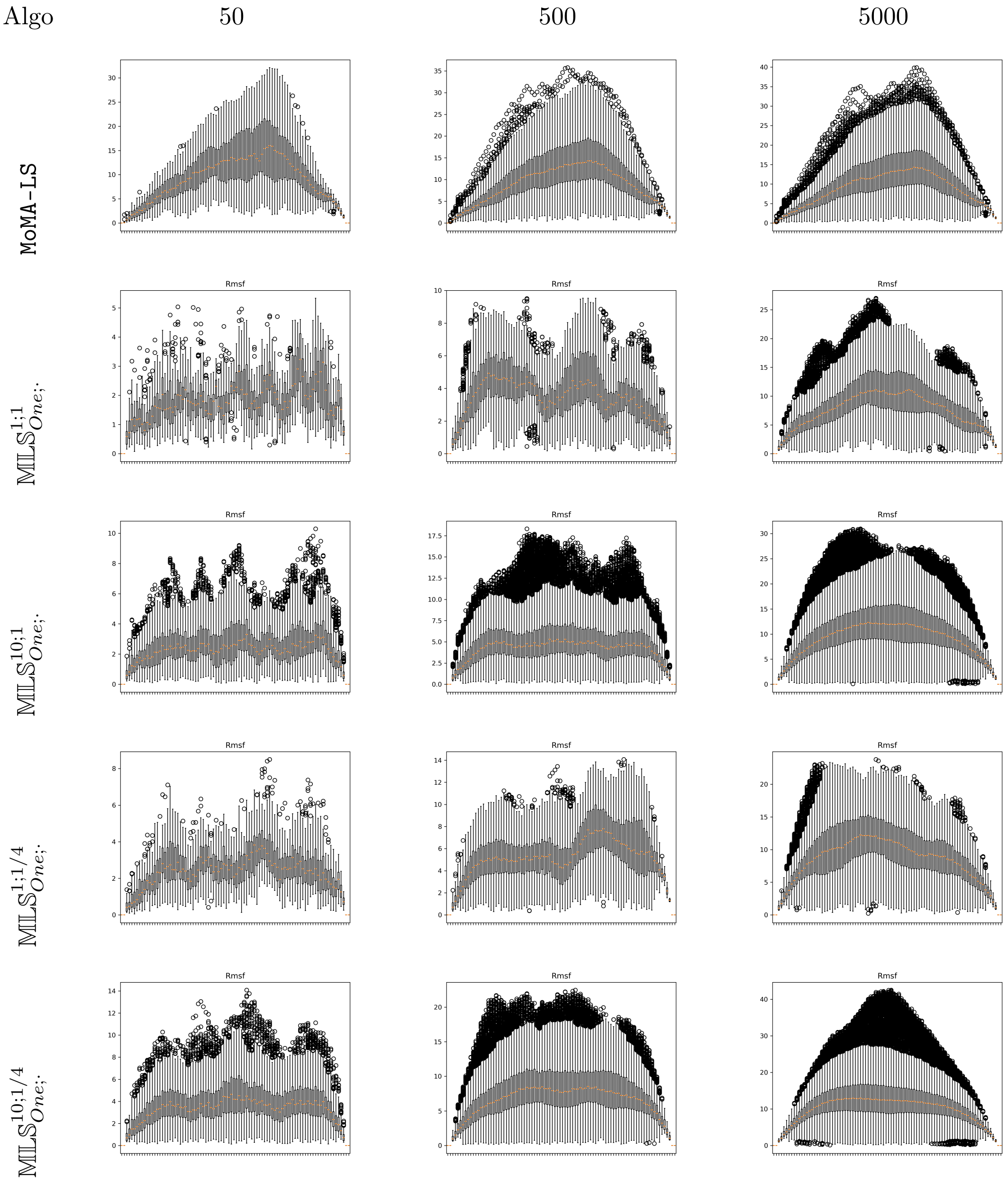


Figure S9: **Loop CDR-H3-HIV: tests with algorithm $\mathbb{MLS}^{N_V;N_{OR}}_{one;N_{ES}}$ and `MoMA-LS`.**

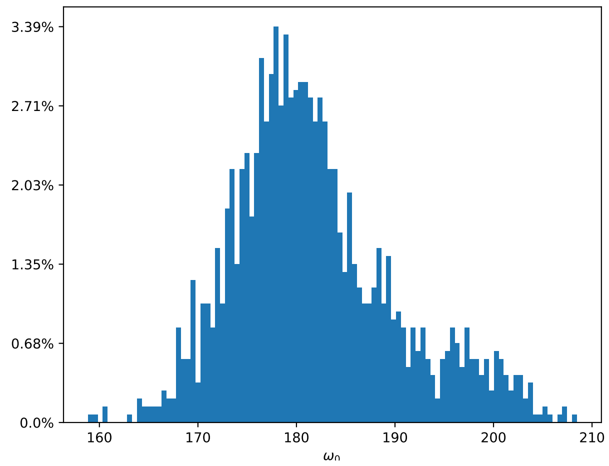


Figure S10: $\omega_0$ **angle values impacting** $C_{\alpha;1}$ **position in** **`MoMA-LS`.** This histogram is made from the sample of 5000 conformations obtained using `MoMA-LS` and $L_0$ of loop PTPN9-MEG2. The $\omega_0$ angle is the torsion angle around the peptide bond preceding the loop.

| | $L_1$ $min/max$lRMSD | $L_2$ $min/max$lRMSD | $L_3$ $min/max$lRMSD |
|---|---|---|---|
| $\mathbf{ULS}^{1;1}_{all;50}[L_0]$ | 0.45/3.14 | 0.44/3.14 | 1.50/3.52 |
| $\mathbf{ULS}^{10;1}_{all;50}[L_0]$ | 0.38/4.37 | 0.39/4.39 | 1.35/4.53 |
| $\mathbf{ULS}^{1;1/4}_{all;50}[L_0]$ | 0.78/3.16 | 0.76/3.16 | 1.56/3.57 |
| $\mathbf{ULS}^{10;1/4}_{all;50}[L_0]$ | 0.45/4.40 | 0.45/4.42 | 1.35/4.73 |
| $\mathbf{ULS}^{1;1}_{all;500}[L_0]$ | 0.45/3.61 | 0.44/3.62 | 1.50/3.83 |
| $\mathbf{ULS}^{10;1}_{all;500}[L_0]$ | 0.38/5.18 | 0.39/5.21 | 1.35/5.05 |
| $\mathbf{ULS}^{1;1/4}_{all;500}[L_0]$ | 0.78/4.74 | 0.76/4.77 | 1.56/5.19 |
| $\mathbf{ULS}^{10;1/4}_{all;500}[L_0]$ | 0.45/5.29 | 0.45/5.33 | 1.35/5.43 |
| $\mathbf{ULS}^{1;1}_{all;5000}[L_0]$ | 0.45/5.27 | 0.44/5.30 | 1.50/5.44 |
| $\mathbf{ULS}^{10;1}_{all;5000}[L_0]$ | 0.38/5.34 | 0.39/5.38 | 1.35/5.66 |
| $\mathbf{ULS}^{1;1/4}_{all;5000}[L_0]$ | 0.78/5.35 | 0.76/5.38 | 1.56/5.53 |
| $\mathbf{ULS}^{10;1/4}_{all;5000}[L_0]$ | 0.45/5.39 | 0.45/5.42 | 1.35/5.85 |
| $\mathbb{MLS}^{1;1}_{all;50}[L_0]$ | 0.41/4.77 | 0.42/4.79 | 1.54/4.50 |
| $\mathbb{MLS}^{10;1}_{all;50}[L_0]$ | 0.37/4.77 | 0.37/4.79 | 1.44/4.54 |
| $\mathbb{MLS}^{1;1/4}_{all;50}[L_0]$ | 1.65/5.30 | 1.63/5.32 | 1.76/5.13 |
| $\mathbb{MLS}^{10;1/4}_{all;50}[L_0]$ | 0.70/5.45 | 0.69/5.46 | 1.54/5.90 |
| $\mathbb{MLS}^{1;1}_{all;500}[L_0]$ | 0.41/5.30 | 0.42/5.32 | 1.54/5.41 |
| $\mathbb{MLS}^{10;1}_{all;500}[L_0]$ | 0.37/5.87 | 0.37/5.90 | 1.39/6.32 |
| $\mathbb{MLS}^{1;1/4}_{all;500}[L_0]$ | 1.55/5.77 | 1.55/5.80 | 1.66/5.94 |
| $\mathbb{MLS}^{10;1/4}_{all;500}[L_0]$ | 0.70/5.95 | 0.69/5.98 | 1.01/6.40 |
| $\mathbb{MLS}^{1;1}_{all;5000}[L_0]$ | 0.41/5.92 | 0.42/5.97 | 1.54/6.17 |
| $\mathbb{MLS}^{10;1}_{all;5000}[L_0]$ | 0.37/6.11 | 0.37/6.14 | 1.01/6.51 |
| $\mathbb{MLS}^{1;1/4}_{all;5000}[L_0]$ | 1.55/5.98 | 1.55/6.00 | 1.51/6.60 |
| $\mathbb{MLS}^{10;1/4}_{all;5000}[L_0]$ | 0.70/6.09 | 0.69/6.12 | 1.01/6.67 |

Table S2: **Loop PTPN9-MEG2: exploration to reach landmark conformations. Using option *All* to retain all solutions per step.** Four conformations of loop PTPN9-MEG2 form two clusters: $L_0, L_1, L_2$ and $L_3$. For `MoMA-LS`, we compute min and max lRMSD distances to these landmarks. For $\mathbf{ULS}^{N_V;N_{OR}}_{\text{One}|\text{All};N_{ES}}$ and $\mathbb{MLS}^{N_V;N_{OR}}_{\text{One}|\text{All};N_{ES}}$, starting from $L_0$, we compute the $max$lRMSD (resp. $min$lRMSD values) values to assess the ability to get away from the cluster (resp. approach conformation $L_3$).
